# Nationwide prediction of type 2 diabetes comorbidities

**DOI:** 10.1101/664722

**Authors:** Piotr Dworzynski, Martin Aasbrenn, Klaus Rostgaard, Mads Melbye, Thomas Alexander Gerds, Henrik Hjalgrim, Tune H Pers

## Abstract

Identification of individuals at risk of developing disease comorbidities represents an important task in tackling the growing personal and societal burdens associated with chronic diseases. We employed machine learning techniques to investigate to what extent data from longitudinal, nationwide Danish health registers can be used to predict individuals at high risk of developing type 2 diabetes (T2D) comorbidities. Based on register data spanning hospitalizations, drug prescriptions and contacts with primary care contractors from >200,000 individuals newly diagnosed with T2D, we used logistic regression-, random forest- and gradient boosting models to predict five-year risk of heart failure (HF), myocardial infarction (MI), stroke (ST), cardiovascular disease (CVD) and chronic kidney disease (CKD). For HF, MI, CVD, and CKD, register-based models outperformed a reference model leveraging canonical individual characteristics by achieving area under the receiver operating characteristic curve improvements of 0.06, 0.03, 0.06, and 0.07, respectively. The top 1,000 patients predicted to be at highest risk exhibited observed incidence ratios exceeding 4.99, 3.52, 2.92 and 4.71 respectively. In summary, prediction of T2D comorbidities utilizing Danish registers led to consistent albeit modest performance improvements over reference models, suggesting that register data could be leveraged to systematically identify individuals at risk of developing disease comorbidities.

## Introduction

Comorbidities of type 2 diabetes (T2D) represent a major cause of death and disabilities resulting in substantial societal and economic burdens^1, 2^. Early interventions have been shown to delay the onset of comorbidities in chronic disease, for instance, intensified multifactorial intervention has been successful in lowering risk of cardiovascular events and slowing down progression of renal disease in patients diagnosed with T2D and microalbuminuria^3^. However, due to the multifactorial underpinnings of chronic diseases and their high prevalence, population-wide interventions remain challenging and hampered by cost-effectiveness and safety concerns^4, 5^. Recent studies^4–7^ and health policy recommendations^2, 8^ suggest that these challenges could be addressed by tailoring interventions to high-risk individuals. Thus, identifying the sub-populations which are at the highest risk of developing chronic disease comorbidities could pave the way to better resources allocation and health outcomes.

Machine learning (ML) techniques are increasingly being used to analyze electronic health record data to predict future disease onset or its future course^9–13^. These efforts include prediction of onset and complications of cardiovascular disease^14–21^, onset of T2D^22–26^, onset of kidney disease^27^, as well as prediction of postoperative outcomes^28–32^, birth related outcomes^33, 34^, mortality^15, 35, 36^ and hospital readmissions^37–43^. However, current approaches typically suffer from a number of limitations. Firstly, although altering the course of chronic disease requires timely prediction^7^, relatively few electronic health record-based studies employ long prediction horizons (a period during which the studies predicted whether a given individual would be diagnosed with a particular outcome). For example, a recent review^9^ noted that out of 81 relevant studies, only 32 used at least a one-year prediction horizon and only 15 used at least 5-year prediction horizon. Second, the above-referenced studies typically do not explicitly test whether the trained prediction model can predict future events^7^. As both medical and data aggregation practices change over time, models trained on data covering a relatively long time period are susceptible to learning patterns, which become irrelevant^44, 45^ – meaning that a model can perform poorly on new data despite having a good average performance on the source data. Lastly, the use of selected cohorts that are not representative of the general population can have important limitations. By relying on data from a specific place of care, state or electronic health record system, the ML model can develop biases specific to that data^7, 9^ and generalize poorly when used for other populations. Overcoming these limitations requires not only an appropriate methodological approach but, importantly, a dataset that is informative, to allow for accurate prediction; long-term, to enable a sufficient prediction horizon and modelling of temporal trends in data; and generalizable, to ensure that the population on which the model is trained is sufficiently representative of the population on which the model is to be applied on.

Most countries do not maintain centralized, nationwide health registers suitable for electronic health record analyses. A notable exception is Denmark, which stores national longitudinal health register data including population-wide information on drug prescriptions, hospital diagnoses, hospital medical procedures as well as claims information from primary care contractors of which all, besides the latter, use international code classifications^46, 47^. For example, there are 348,000 individuals diagnosed with T2D from 1995 to 2011. For these individuals, the registers contain information on >14.2 million hospital diagnoses (>15,900 unique codes), >21.3 million procedures (>13,000 unique codes), >182 million claims from primary care contractors (>12,600 unique) and >143.7 millions prescriptions (22,000 unique) amounting to a total of >553.2 million medical events. In sum, the Danish longitudinal registers contain the approximate medical life course of virtually every Danish citizen, possibly presenting valuable opportunities for predicting adverse medical outcomes, such as comorbidities in the chronically ill.

In this study, we leveraged Danish register data and commonly used ML methods to assess the extent to which nationwide longitudinal records on hospital diagnoses, hospital procedures, prescriptions and claims from primary care contractors covering 10 years of nearly >200,000 T2D patients, could predict future onset of chronic disease comorbidities in these patients. We chose to focus on T2D due to its prevalent nature, its diverse commodities and because early intervention has been shown to reduce subsequent risk of developing commodities. Applying three common ML methods, namely logistic ridge regression, random forest and gradient boosting on the above-mentioned comprehensive Danish population-wide register data we predicted future onset of MI, HF, CVD, CKD, stroke and all-cause mortality within a five-year window of individuals’ individuals’ first T2D diagnosis.

## Materials and methods

### Danish population-wide registers

The Danish National Patient Register^48^ contains information on all non-psychiatric hospital visits since 1977 and includes psychiatric hospital, emergency department and outpatient clinic visits since 1995. All medical events are recorded using Health Care Classification System which combines multiple Danish and international classifications. Notably, all diagnoses are recorded using the International Statistical Classification of Diseases and Related Health Problems (ICD) system (revision 8 used between 1977 and 1994 and revision 10 used since 1995); surgeries from 1996 onwards are recorded using the Nordic Medico-Statistical Committee Classification of Surgical Procedures (NCSP); treatment and diagnostic procedures are represented through unique Danish hierarchical classifications. The Danish National Prescription Register^49^ contains detailed information on all prescription drugs sold in Danish pharmacies since 1995 and uses the Anatomical Therapeutic Chemical (ATC) classification system. The Danish National Health Service Register contains claims information on “activities of health professionals such as general practitioners, practising medical specialists, physiotherapists, dentists, psychologists, chiropractors, and chiropodists”^50^ and uses a unique classification scheme. Furthermore, we included data from the Central Person Register comprising an individual’s sex, date of birth, country or region of birth, date of death as well as generalized residence address (five areas throughout Denmark). The pregnancy and childbirth information used to identify gestational diabetes was extracted from The Danish Medical Birth Register^51^. Causes of death were identified through the Danish Register of Causes of Death^52^. More detailed information regarding the registers can be found in Table 1.

**Table 1.** Overview of Danish national health registers employed in this study. *ICD-8 used between 1977 and 1995, ICD-10 between 1995 and 2016. ICD-10 stands for 10th revision of International Statistical Classification of Diseases and Related Health Problems; DN, Danish National; NCSP, Nordic Medico-Statistical Committee Classification of Surgical Procedures; ATC, Anatomical Therapeutic Chemical Classification; #, number of.

| Register | Data | Int. classification | Dates | Total #records |
| --- | --- | --- | --- | --- |
| DN Patient Register | Hospital diagnoses | ICD-10 | 1995 → 2016* | 179.0 million |
|  | Surgical procedures | NCSP | 1996 → 2016 | 26.9 million |
|  | Treatment procedures | — | 1999 → 2016 | 103.4 million |
|  | Diagnostic procedures | — | 1999 → 2016 | 62.5 million |
| DN Prescription Register | Drug prescriptions | ATC | 1994 → 2016 | 930.4 million |
| DN Health Service Register | Claims data | — | 1990 → 2016 | 2,147.0 million |
| DN Medical Birth Register | Pregnancies & prenatal care | — | 1973 → 2016 | 2.7 million |

### Study population and inclusion criteria

We employed an observational prediction study using a five-year prediction horizon with a time of prediction set to date of an individual’s first T2D diagnosis. To that end, we used the Danish registers to sample a population of (n=203,517) T2D patients. The date of the first T2D diagnosis was defined as the date of first registration of hospital T2D diagnosis (ICD10 code E11), prescription of insulin and analogues (ATC code A10A) or prescription of blood glucose lowering drugs (ATC code A10B). We excluded A10B prescriptions which occurred during pregnancy as well as individuals who had their first A10A prescription before the age of 30. We required the date of prediction to occur between January 1 2000 and December 31 2010 to prevent inflation of first T2D diagnoses during the first years of register information availability and to establish five-year followup for all patients. Additionally, to evaluate the predictiveness of medical events following the first T2D diagnosis, we also sampled four populations of T2D patients with time of prediction set to one year (n=181,100), two years (n=165,482), three years (n=150,228), and four years (n=137,240) after the first T2D diagnosis (maintaining the five-year prediction horizon).

For each of the five T2D comorbidities, as the prediction objective we chose the five-year risk of an individual’s first diagnosis of a specific comorbidity. Specifically, we chose the first occurrence of a non-referral ICD-10 hospital diagnosis code within the Danish National Patient Register or ICD-10 code assignment as cause of death in the Danish Register of Causes of Death related to HF (ICD-10 code H50), MI (ICD10 code I21), ST (ICD-10 code I61-I64), other CVD (ICD-10 codes I20, I22-25, I679) and CKD (ICD-10 code N17-N19). For each comorbidity, a case was defined as an individual with a first-time diagnosis of the given comorbidity occurring within the five-year prediction horizon, and similarly, a non-case was defined as an individual with no diagnosis of the given comorbidity occurring during the prediction horizon. Consequently, individuals who died during the prediction horizon period without prior comorbidity diagnosis were treated as non-cases. To avoid instances where the comorbidity was diagnosed before the diagnosed onset of T2D (for example through an informative diagnostic test ordered to confirm prior beliefs about a given outcome), we required comorbidities to be diagnosed at least 30 days after the diagnosed date of T2D onset (henceforth *buffer period*). In addition to predicting T2D comorbidities, we also predicted all-cause mortality defining cases and non-cases based on the Central Person Register and the Danish Register of Causes of Death. For a study overview and study population characteristics, please refer to Table 2, Fig. 1 and Supplementary Table 1.

**Table 2.**
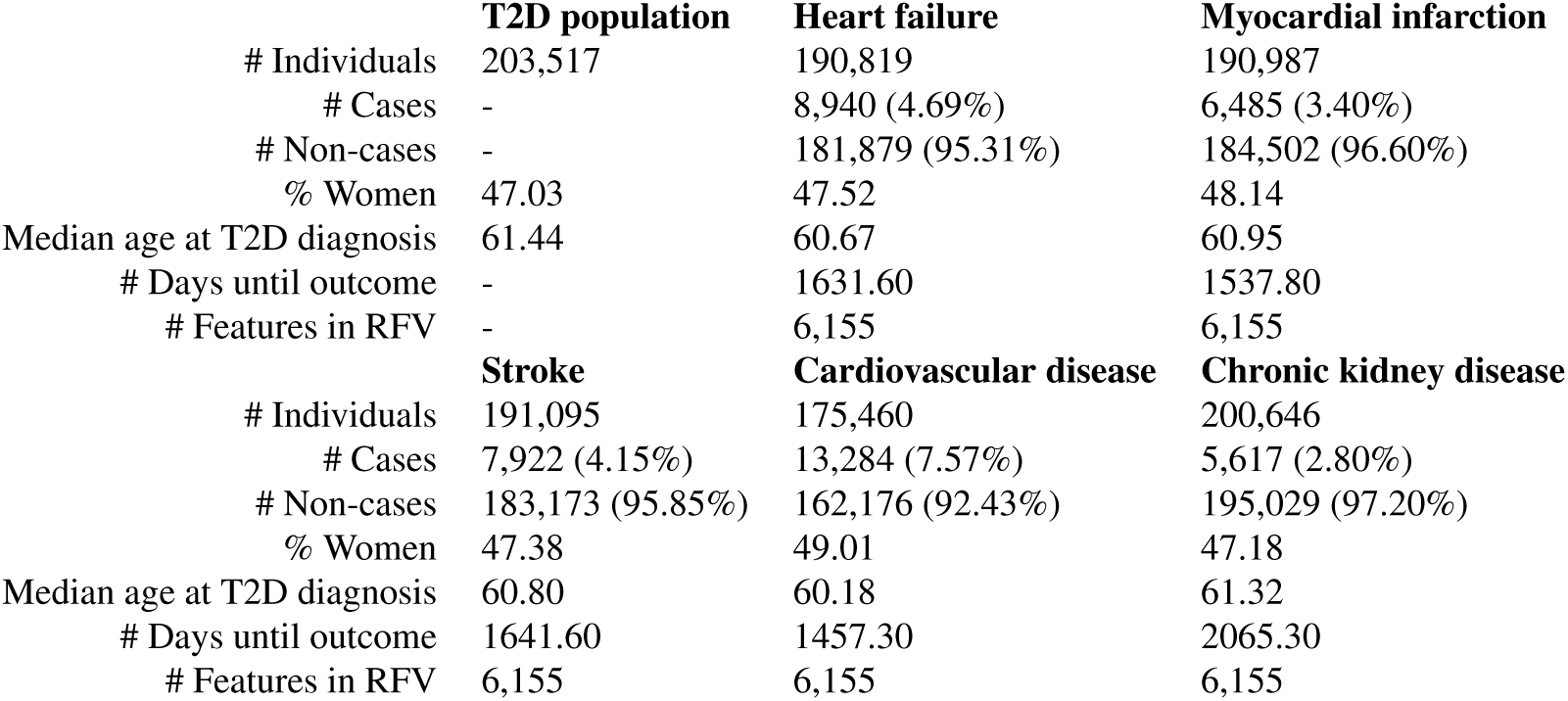
Overview of study population characteristics among all newly diagnosed type 2 diabetics (T2D population) and five comorbidity populations (individuals free of specific comorbidity at first T2D diagnosis). RFV, register feature vector; #, number of.

|  | T2D population | Heart failure | Myocardial infarction |
| --- | --- | --- | --- |
| # Individuals | 203,517 | 190,819 | 190,987 |
| # Cases | - | 8,940 (4.69%) | 6,485 (3.40%) |
| # Non-cases | - | 181,879 (95.31%) | 184,502 (96.60%) |
| % Women | 47.03 | 47.52 | 48.14 |
| Median age at T2D diagnosis | 61.44 | 60.67 | 60.95 |
| # Days until outcome | - | 1631.60 | 1537.80 |
| # Features in RFV | - | 6,155 | 6,155 |
|  | Stroke | Cardiovascular disease | Chronic kidney disease |
| # Individuals | 191,095 | 175,460 | 200,646 |
| # Cases | 7,922 (4.15%) | 13,284 (7.57%) | 5,617 (2.80%) |
| # Non-cases | 183,173 (95.85%) | 162,176 (92.43%) | 195,029 (97.20%) |
| % Women | 47.38 | 49.01 | 47.18 |
| Median age at T2D diagnosis | 60.80 | 60.18 | 61.32 |
| # Days until outcome | 1641.60 | 1457.30 | 2065.30 |
| # Features in RFV | 6,155 | 6,155 | 6,155 |

**Figure 1.**
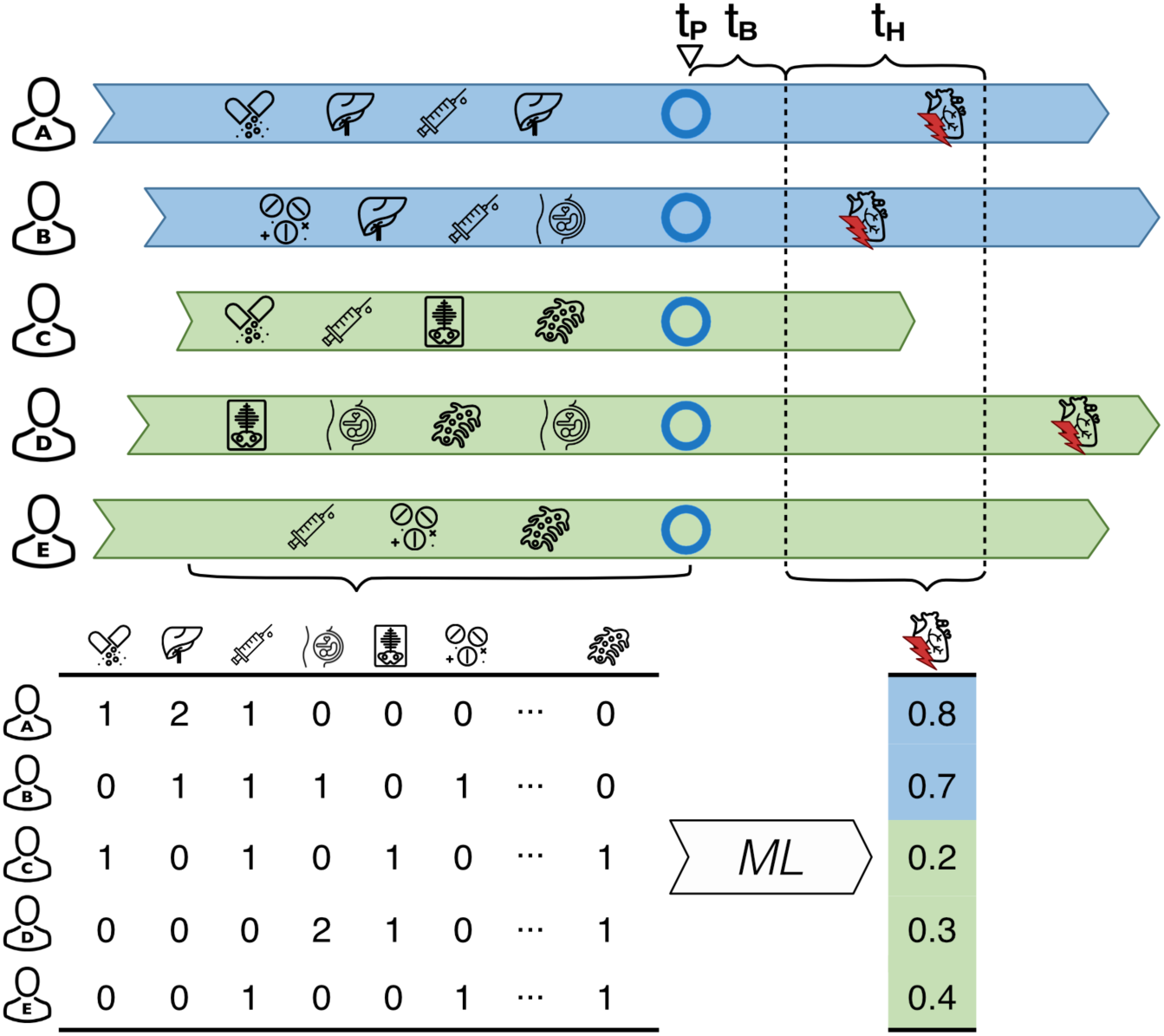
The colored arrows represent each individual’s accumulated register events. *t*_*P*_ represents the date of prediction (in this case the day of the first T2D diagnosis represented by blue circles). *t*_*B*_ depicts the buffer period of 30 days used to exclude individuals who were diagnosed with the comorbidity shortly before *t*_*P*_. *t_H_* represents the five-year prediction horizon. Individuals for whom the comorbidity occurred before the prediction horizon were removed. Individuals with their first comorbidity diagnosis occurring within the prediction horizon are referred to as *cases* and all others are referred to as *non-cases*. For each individual, register events before the time of prediction are aggregated into counts (table below) and used as the prediction model’s input vector to determine the likelihoods of being a case or a non-case. ML, machine learning.

### Predictor variables

For all individuals, we created two sets of predictive variables (henceforth *feature vectors*): a canonical feature vector, composed of canonical features, to be used by a reference model, and a register feature vector, composed of canonical features as well as features extracted from Danish health registers, to be used by the ML models. The canonical features were sex (binary encoded), country or region of birth (as denoted in the Central Person Register, 13 categories), date of prediction (continuous), age at date of prediction (continuous), number of days lived in a general region of Denmark (continuous, five regions), an interaction term between age and sex, a restricted cubic spline of the prediction date and a restricted cubic spline of age at date of prediction. The register features were counts of individuals’ hospital diagnoses, drug prescriptions, procedures and interactions with primary care contractors. Diagnoses were limited to only those that were non-referral and encoded in the ICD-10 system, while information on claims from primary care contractors was encoded as an individual’s count of interactions with a specific health care contractor type (e.g. “general practitioner” or “dentist”). Each diagnosis, drug prescription, and medical procedure was included at multiple levels of specificity by leveraging their respective code hierarchies (ICD-10 for diagnoses, NCSP for procedures and ATC for drug prescriptions). For example, the diagnosis “other type of myocardial infarction” (ICD-10 code I21.A) was represented by three features, namely diagnosis of “diseases of the circulatory system” (ICD-10 code I), diagnosis of “acute myocardial infarction” (ICD-10 code I21) and diagnosis of “other type of myocardial infarction” (ICD-10 code I21.A). The value of each feature for a given individual was the cumulative number of observations of electronic health record codes representing that feature (across multiple diagnoses or prescriptions) prior to the time of prediction. The register feature vector for all comorbidities comprised 6,181 register features: 26 canonical features, 3,423 hospital diagnoses, 2,015 hospital procedures, 670 prescriptions and 47 primary care interactions.

### Prediction models

For each comorbidity (and all-cause mortality), four algorithms were used to predict an individual’s risk of developing the comorbidity within the prediction horizon. Logistic ridge regression^53^ with the canonical feature vector as input was used as the reference model. Restricted cubic splines were used to model non-linear effects of age separately for the two genders by adding an interaction term. As our register-based models, we used three algorithms (model types), namely logistic ridge regression, random forest^54^, and decision tree gradient boosting^55^ with the register feature vector as input. Briefly, logistic ridge regression is a logistic regression model where parameter estimates are shrunk towards zero by means of a penalty term (tuning: penalty parameter, maximum number of iterations); random forest is an ensemble model which in our case consists of many classification trees built in many bootstrap samples (tuning: number of trees, maximum depth of each tree); decision tree gradient boosting results in a sequence of shallow decision trees where each consecutive tree is trained to improve the performance of the ones before it (tuning: learning rate, maximum depth of each tree).

### Dataset split and model training/evaluation procedures

For each comorbidity (and all-cause mortality), each prediction model was trained and evaluated using the following procedure. First, all individuals were split into three sets, namely a training sample, a test sample, and a validation sample. The training set was allocated the first 70% of the individuals (sorted by date of first T2D diagnosis) of the full dataset and used for three-fold cross-validated^56^ hyperparameter search. Similarly to work by Corey *et al.*^28^, the remaining (most recently diagnosed) 30% of the full dataset were randomly split into two balanced (the proportion of cases was maintained in each split) subsets: a test set which was allocated 20% of the full dataset to be used for model selection for each outcome and the validation set which was allocated the remaining 10% of the full dataset and exclusively used to present final results. For each model type, the best parameter set was chosen by first repeatedly training the model on two-thirds of the training data and evaluating it on the remaining third, and then averaging accuracy measures across the three runs. The parameterization resulting in the highest average area under the receiver operating characteristic curve (AUROC) was used to re-train the model on the combined training set yielding the best model for a given model type. The best model’s performance was evaluated against the test set and compared to the corresponding best models from other model types. Finally, the validation set was used to report the best models performance (reporting AUROC). 95% confidence intervals of AUROC and AUROC differences (DAUROC) between models were obtained using the bootstrap method^57^ using 1,000 samples. The training and validation procedure is illustrated in Figure 2 and an overview of all tested parameters can be found in Supplementary Table 6. The analysis was implemented in Python programming language^58^ version 3.6.2 using modules numpy^59^ version 1.13.1, pandas^60^ version 0.20.3, scikit-learn^61^ version 0.19.0, patsy^62^ version 0.4.1, xgboost^63^ version 0.80 and plots were made using bokeh visualization library^64^ version 1.0.4.

**Figure 2.**
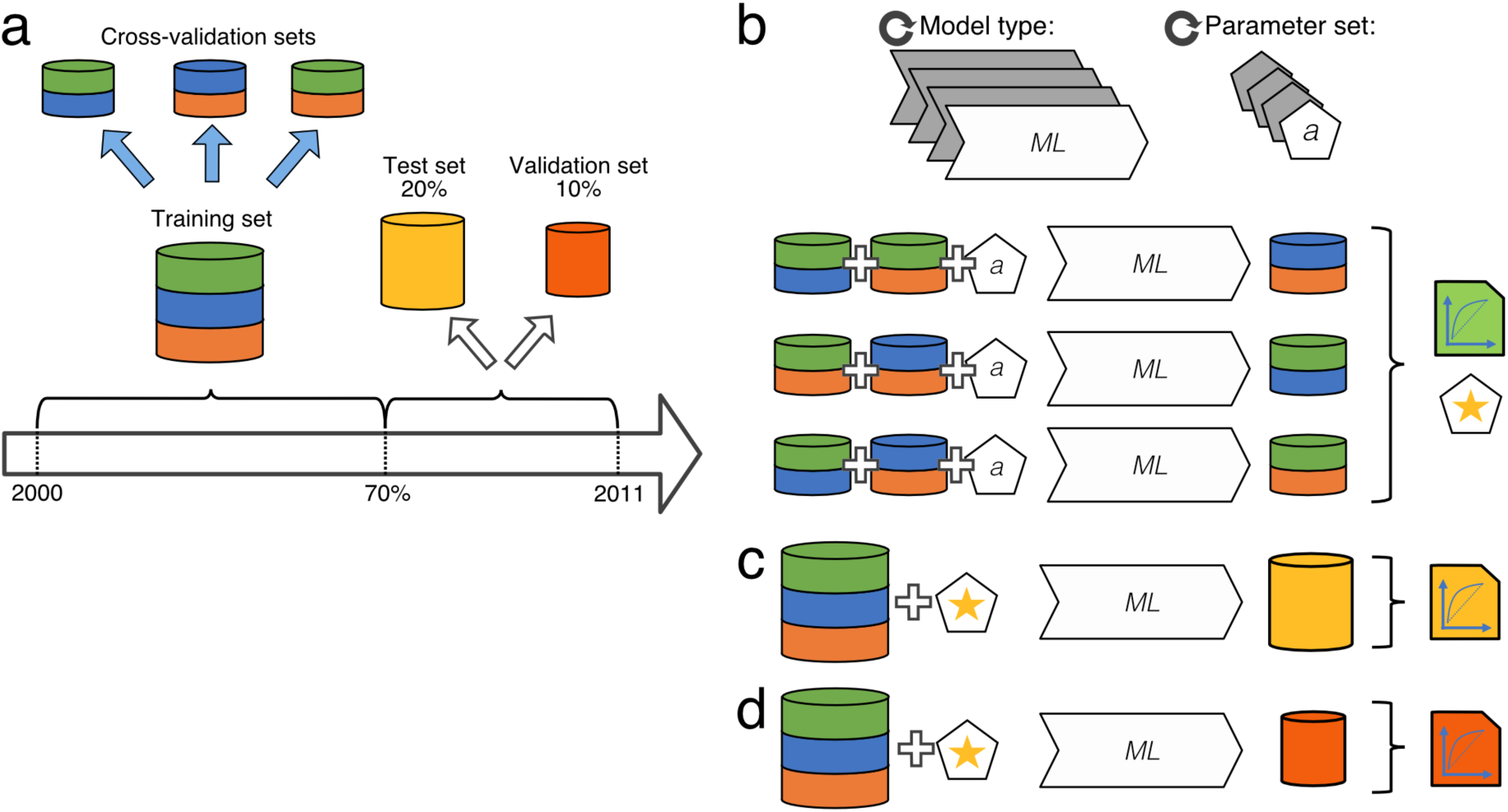
Data split and model tuning. (**a**) Data were split so that the training set constituted the first 70% of the data (time-wise, according to the time of the first T2D diagnosis). The test and validation sets were divided by balanced random sampling from the remainder (time-wise latter part) of the dataset. (**b**) For each model type, the best parameter set was chosen through its evaluation following three-fold cross-validation on the training data. (**c**) For each model type, the best model was obtained by re-training the model on the entirety of the training set using the best parameter set. Performance of each best model was evaluated on the test set for the purpose of development of this work. (**d**) Performance of each best model was evaluated on the validation set for the purpose of reporting the results. ML, machine learning.

## Results

### Prediction of T2D comorbidities based on clinical events incurred prior to the first T2D diagnosis

To assess to what extent register data can predict comorbidities for T2D patients, we first identified all individuals either diagnosed with T2D or being prescribed diabetes drugs between 2000 and 2011 (See Material and Methods; Fig. 1). Among these 203,517 individuals, we next identified subsets having received a hospital diagnosis of a major T2D comorbidity within a five-year period following their first T2D diagnosis. We identified 8,940 incident cases for HF, 6,485 for MI, 7,922 for ST, 13,284 for CVD and 5,617 for CKD, and treated the remaining individuals as non-cases (see Table 2 for study population characteristics).

The average number of clinical events used for the prediction per individual was 737 for HF, 759 for MI, 761 for ST, 697 for CVD, 776 for CKD. For example, for CVD, each individual had an average of 41.9 hospital diagnosis events (the most common being essential hypertension, ICD10 code I109), 111.8 hospital procedures (the most common being x-ray examination, SKS code UXR), 6.79 claims data events (the most common being contact with general practitioner, SSR category 80), 536.9 drug prescriptions (the most common being antibacterials for systemic use, ATC code J01) prior to their first T2D diagnosis.

Using the AUROC measure, we first assessed how well we could predict the five T2D comorbidities based on clinical events incurred prior to the individuals’ first T2D diagnosis. The gradient boosting model yielded the highest AUROC for all outcomes and outperformed the reference model for four comorbidities (HF, MI, CVD, CKD; Table 3). The best gradient boosting models performed better than the best random forest models for all comorbidities and logistic ridge regression for HF, CVD and CKD. Additionally, register-based logistic ridge regression outperformed the reference model for HF, MI, CVD, and CKD. Together these results indicate that prediction based on hundreds of features assembled across health registers improves on prediction based on a limited number of canonical measures typically used to assess risk. Moreover, our results suggest that gradient boosting improves on logistic ridge regression models’ performance in the context of large-scale health register data-based prediction. The following analyses were focused on predicting the HF, MI, CVD and CKD comorbidities (see Supplementary Fig. 2 for results on ST). Detailed results for each tested model parametrization can be found in Supplementary Table 2 and 3.

**Table 3.**
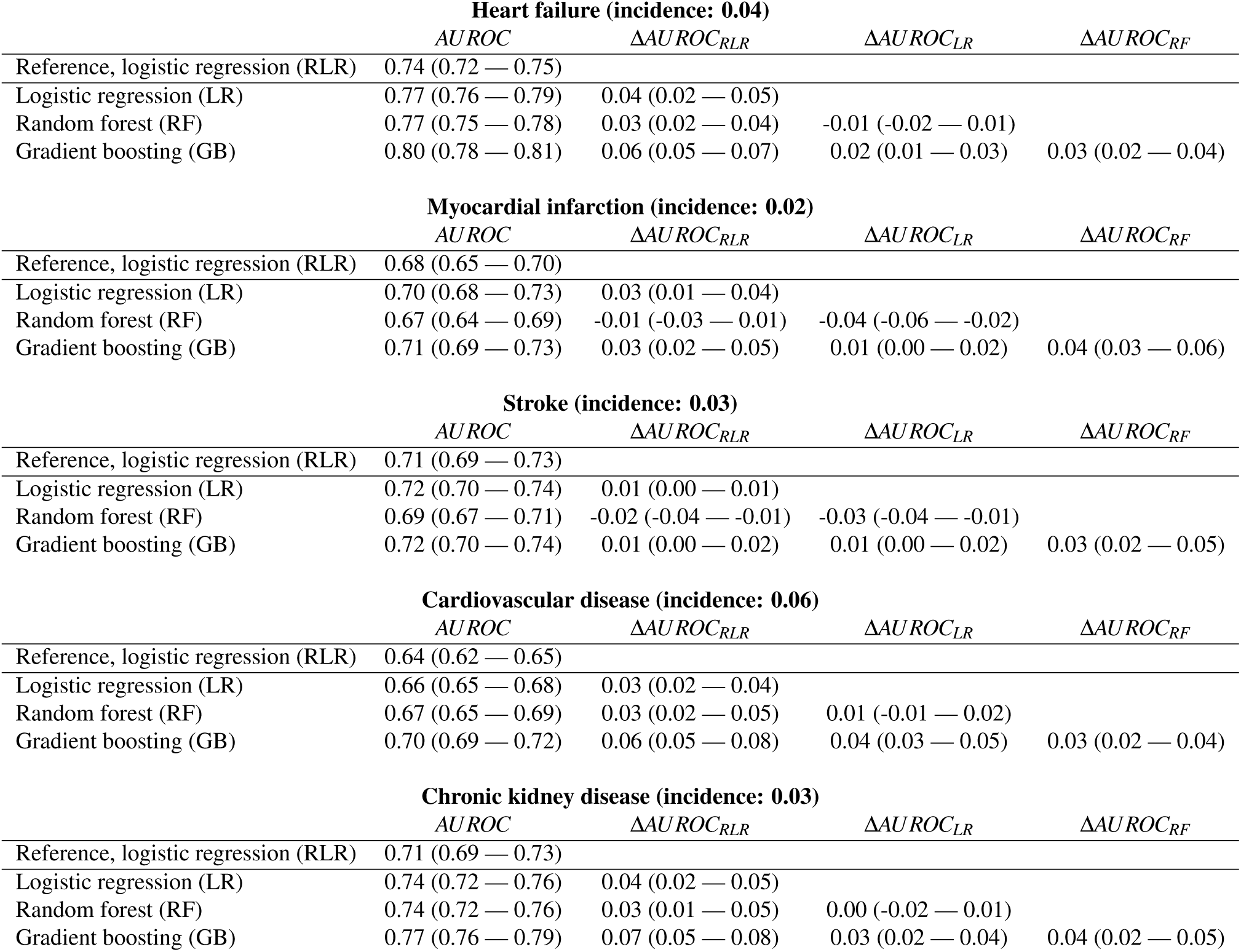
AUROC measures for each prediction model’s best parameterization. We applied a reference- and three register-based models on fifteen years of health register data comprising hospital diagnoses, hospital procedures, drug prescriptions and interactions with primary care contractors to predict five-year risk for the five T2D comorbidities. For each comorbidity, prediction was performed on a T2D population free of that comorbidity at the date of prediction (date of individuals’ first T2D diagnosis). The reference model was a logistic ridge regression based on canonical features: age, sex, country or region of birth and date of first T2D diagnosis as well as their interactions, while the register-based models were logistic ridge regression, random forest and gradient boosting based on the canonical features as well as hospital diagnoses, hospital procedures, drug prescriptions and interactions with primary care extracted from Danish health registers. Incidences are proportions of cases within comorbidities’ sub-population at the end of the prediction horizon. Value ranges in brackets represent 95% confidence intervals based on bootstrap sampling. For heart failure, myocardial infarction, cardiovascular disease and chronic kidney disease the gradient boosting model outperformed the reference models. AUROC, area under receiver operating characteristic curve.

### Comorbidity incidences across predicted risk percentiles

To investigate the models’ ability to rank individuals according to their risk of being diagnosed with a given comorbidity, we, for each comorbidity, grouped individuals into risk percentiles (based on the predicted risk from the gradient boosting and reference models) and plotted for each percentile the five-year comorbidity incidence (i.e. the fraction of individuals at that percentile incurring a given comorbidity during the prediction horizon; Fig. 3). Individuals in the highest percentiles for both the reference and register-based models exhibited markedly increased comorbidity incidences compared to the rest of the population. For example, for the gradient boosting model, individuals predicted to be in the 95th percentile had incidence risk ratios of 3.67 for HF, 2.66 for MI, 3.11 for CVD and 4.18 for CKD. Individuals predicted to be in the top gradient boosting-based risk percentiles had modest, but consistently higher five-year comorbidity incidence compared to individuals grouped by the reference model into corresponding top percentiles. This finding underscores that prediction based on multiple register-derived events improved on models including a more limited number of canonical features.

**Figure 3.**
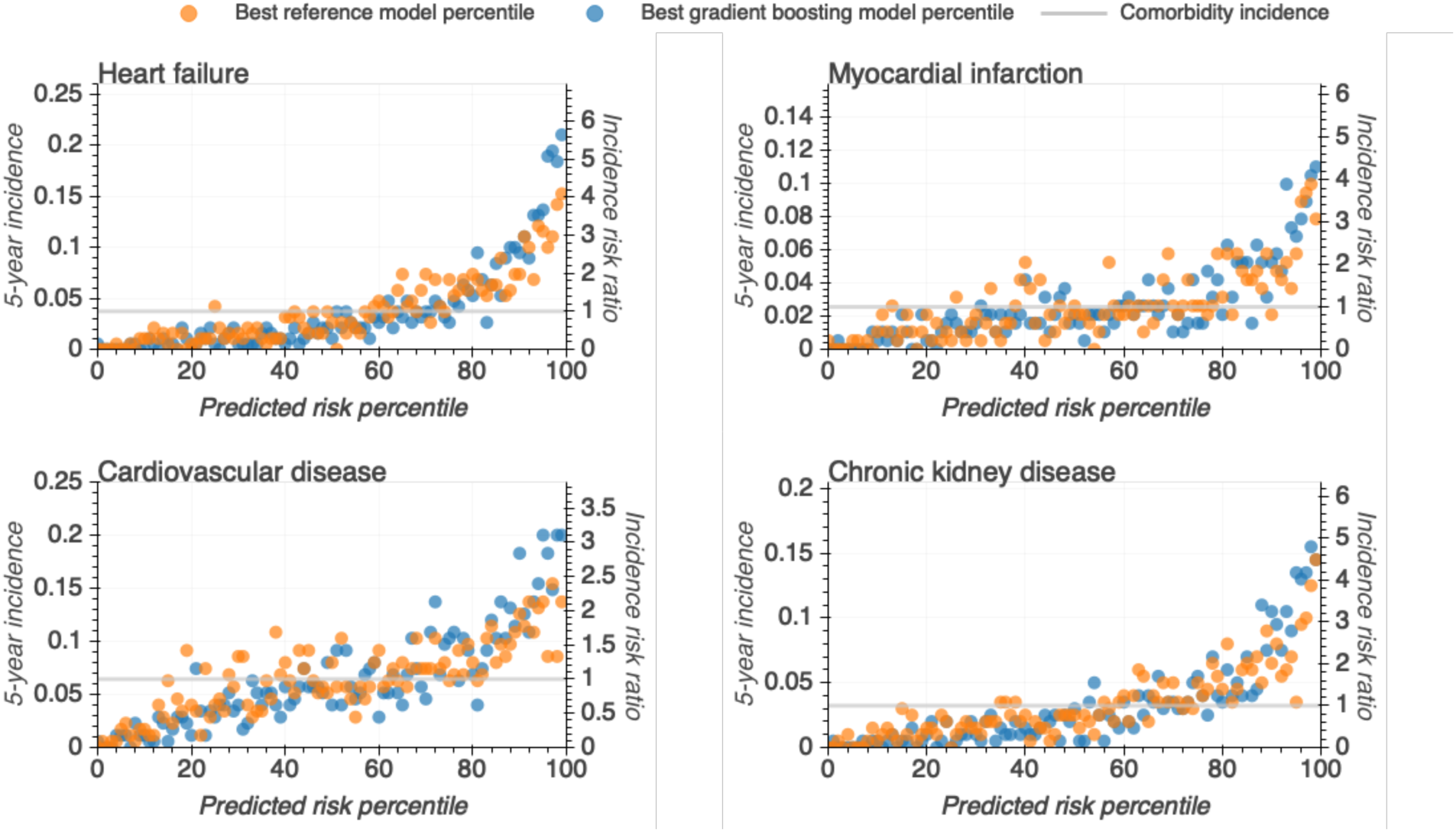
All individuals were ranked according to their predicted risk (in increasing order) by the best gradient boosting (blue) and best reference (orange) models and binned into percentiles. Plotted are the observed five-year comorbidity incidences for individuals in each percentile. Left y-axis; incidence defined as the observed proportion of individuals who did develop the given comorbidity within the five-year prediction horizon. Right y-axis; the incidence risk ratio defined as a ratio between the percentiles’ and population observed five-year comorbidity incidence. Gray horizontal line; population five-year comorbidity incidence.

A potential application of population-wide risk prediction models is to identify individuals at risk of a certain comorbidity (or another outcome). To further investigate whether individuals predicted to be at high-risk in fact tended to be diagnosed with the given comorbidity, we computed, at different thresholds at the right tail of the predicted risk distributions, five-year comorbidity incidences for the best gradient boosting and reference models. At each threshold, we calculated a risk ratio, by comparing the five-year incidence of each of the four comorbidities for all individuals above that threshold (e.g. the proportion of cases to non-cases among the 1,000 most at-risk predicted individuals) to the T2D patient population-wide incidence. For the gradient boosting model of HF, CVD and CKD, we observed that the risk ratios were significantly higher than those based on the reference models (confidence intervals widened towards the rightmost end of the distribution due to the decreasing number of individuals for smaller thresholds).

### Predictive features

To gain insight into the decision-making process of the best models’ predictions, we analyzed the gradient boosting models’ feature importance, i.e. the underlying features’ relative contribution to the prediction. First, we assessed the contribution of each feature type (canonical features, prescriptions, address information, hospital diagnoses, and procedures, primary care interactions) for the 50 most predictive features as well as their cumulative importance (sum of all feature importance for each feature type; Fig. 5). We found that feature importance formed a long right-skewed distribution, indicating that model predictions were based on a large number of features. Furthermore, we identified prescription features as being the most important for all comorbidities (feature importance of 30% for ST to 41% for HF), followed by either canonical features (17% for HF to 27% for ST) or hospital diagnoses (20% for CKD to 29% for CVD). Detailed information on the first seven most important features for each outcome as well as their distribution in cases and non-cases populations can be found in Supplementary Fig. 3. Together, these results show that data from health registers, especially prescriptions and hospital diagnoses, provide predictive information on top of canonical features such as age, sex, and date of diagnosis, which are known to be predictive for the given comorbidities.

**Figure 4.**
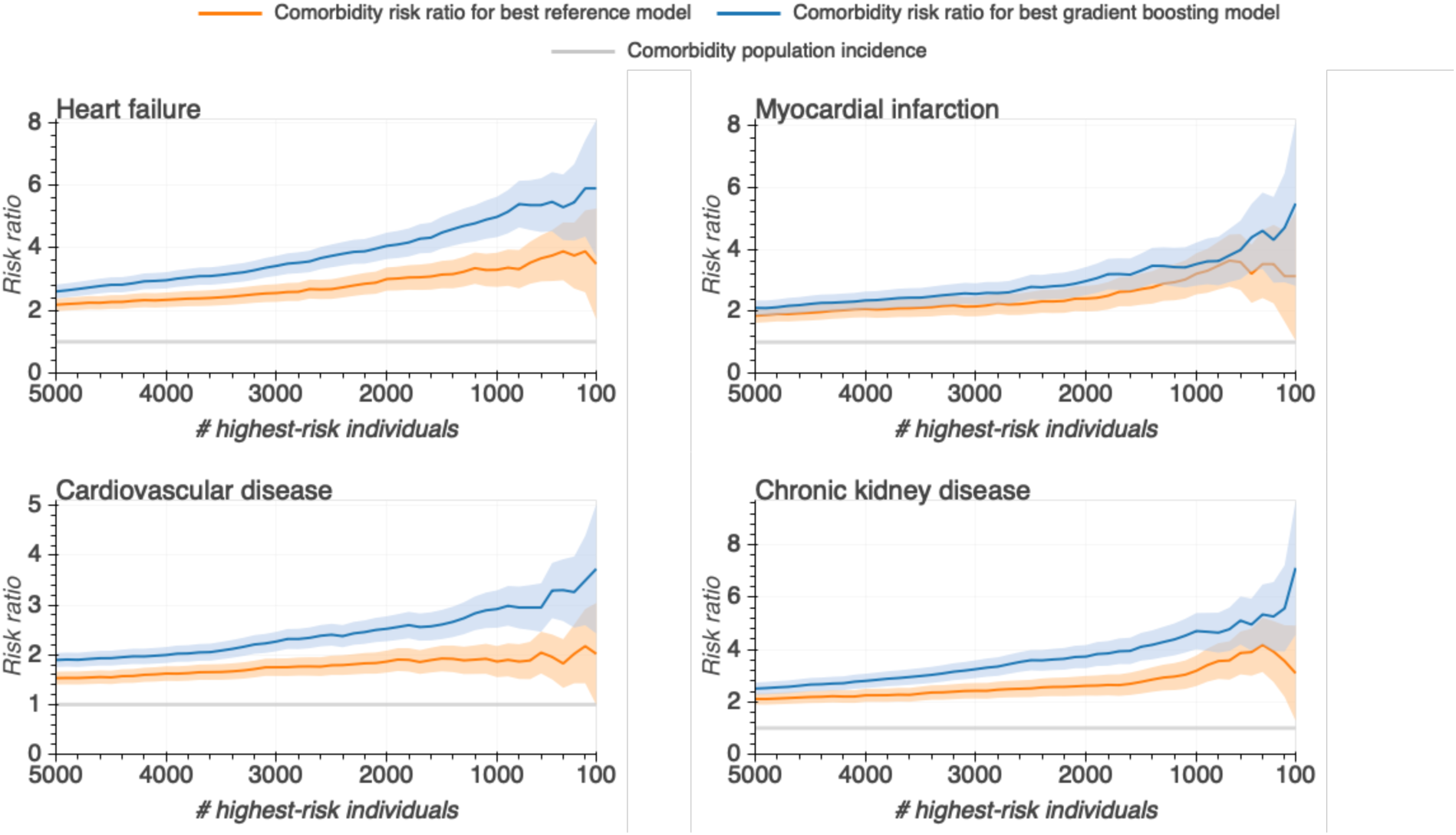
For each comorbidity individuals were ranked according to their predicted risk by the gradient boosting (blue) and reference (orange) models. For a number of individuals predicted to have the highest risk, risk ratios were calculated as the comorbidity incidence of individuals ranking above that threshold over the comorbidity incidence in the entire study population. 95% confidence intervals (shaded areas) were obtained through bootstrap sampling.

**Figure 5.**
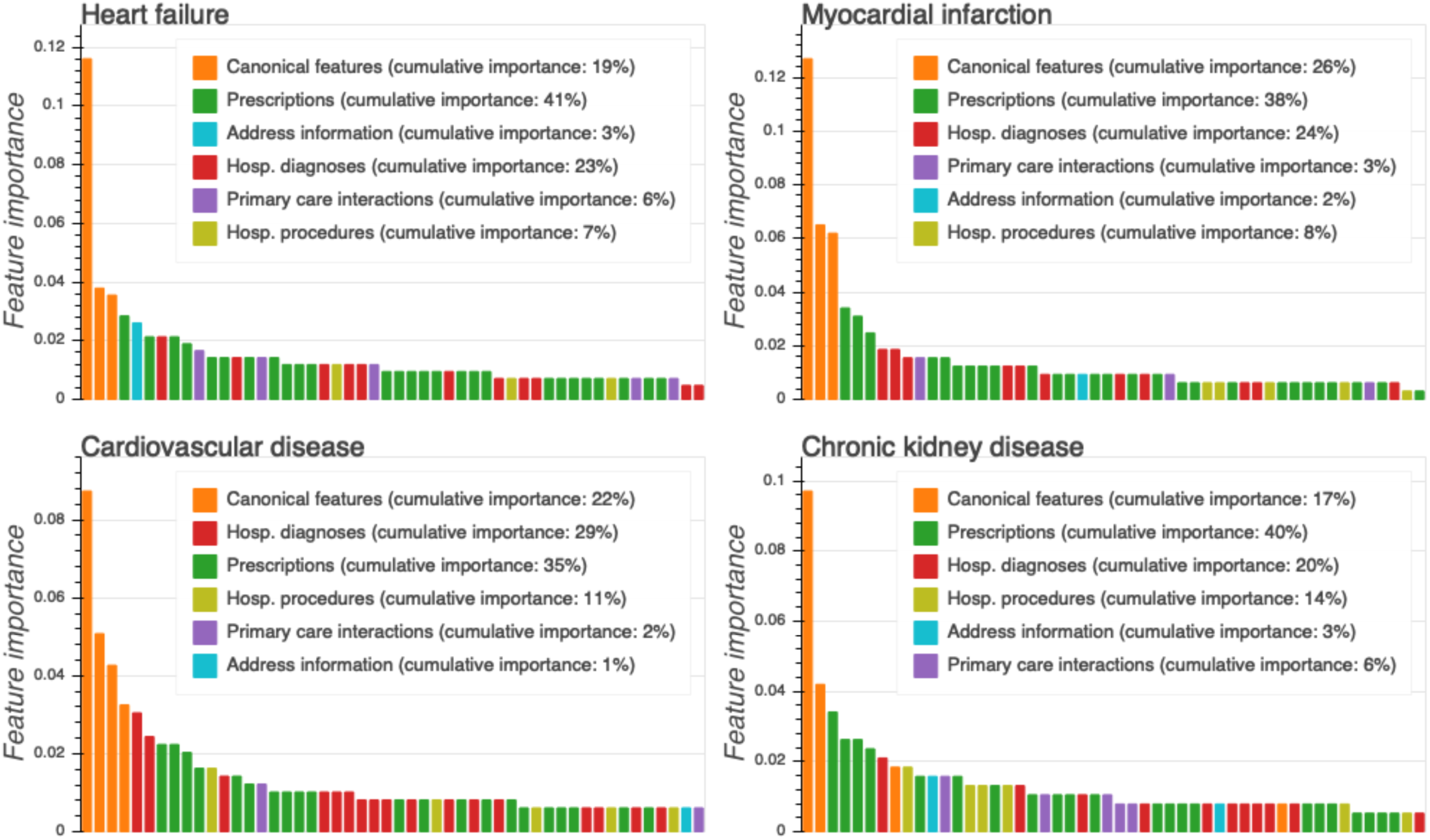
Top 50 most predictive register feature vector features from the best gradient boosting model ranked by their importance and colour-coded according to type (x-axis). Feature importance is a normalized estimate of a relative contribution of the feature to the model prediction (y-axis). Drug prescription features had the highest overall cumulative importance followed by the canonical features and hospital diagnoses. Age, interaction between age and sex, and date of first T2D diagnoses were the three most important features for all comorbidities.

### Inclusion of clinical diagnoses incurred after first T2D diagnosis does not improve prediction

Having shown that the inclusion of large-scale register health data can improve prediction of T2D comorbidities, we next investigated whether the inclusion of clinical events following the first T2D diagnosis could further improve prediction accuracy. Towards this aim, we employed the same prediction procedure with the time of prediction set to one-, two-, three- and four years after the first T2D diagnosis while keeping the length of the buffer period (30 days) and prediction horizon (five years) constant. Contrarily to our expectation, we observed largely unchanged model performances in terms of AUROC (Supplementary Table 5). From a technical standpoint, these results indicate that clinical events succeeding the first T2D diagnosis do not contain new information predictive of the T2D comorbidities. These results suggest that register-based prediction models are equally accurate when predicting T2D comorbidities at the date of first T2D diagnosis and during the following four years. For information on population characteristics and model performances please refer to the Supplementary Tables 1 and 4.

### Comparison between time-split and non-time-split prediction

A commonly criticized aspect of register-based prediction analysis is that models are evaluated on data covering the same time period as the training data^7, 8^. In so doing, it is impossible to determine whether data patterns learned by the models are predictive for future events due to potential changes in medical practice or changes in health patterns in general. In this work, we applied a so-called *time-split* approach (training set comprising of the first 70% of the population based on date of first T2D diagnosis, with the test and validation sets randomly sampled from the remaining 30%). To test whether sampling of the training, test and validation sets without taking into account individuals’ dates of first T2D diagnoses, would yield different results, we re-ran all analysis with such a *non-*time-split approach. Expectedly, the AUROCs from the non-time-split approach were consistently, although non significantly, better than AUROCs from the time-split approach applied here (Supplementary Table 7). These findings suggest that medical practice predictive of comorbidities slightly changed between January 1 2000 and December 31 2010. The consistent differences underline the potential bias of models trained in a non-time-split approach towards overestimating prediction accuracies of future events.

### Death as a competing risk

Individuals who died during the prediction horizon before potentially being diagnosed with the comorbidity were categorized as non-cases. Consequently, death constituted a competing risk to the comorbidity diagnosis. ML models could potentially account for death as a competing risk by implicitly weighting the risk of a given comorbidity diagnosis against the risk of death prior to receiving the given diagnosis. To further explore that issue, we first trained and evaluated prediction performance for all-cause mortality using the same procedure as for T2D comorbidities. We found that gradient boosting significantly outperformed all other models (Table 4) with AUROCs substantially higher compared to the best AUROCs observed for the T2D comorbidities (AUROC=0.87 vs. HF AUROC=0.80). Furthermore, the gradient boosting feature importances for all-cause mortality, in contrast to the comorbidity prediction feature importances, suggested a more prominent role of hospital diagnoses compared to canonical features and prescriptions (cumulative importances of 32%, 19% and 29%, compared to average cumulative importances of 24.2%, 22.2% and 36.8%, respectively; Fig. 6c). Among the top seven predicted features were, expectedly, age and sex but also three types of cancer diagnoses (malignant neoplasm of breast, pancreas, brain; Supplementary Fig. 3). For each comorbidity and the gradient boosting and the reference model, we next investigated five-year incidence of all-cause mortality in each population percentile (in the distribution derived by ranking individuals based on their predicted risk of a given comorbidity). For comorbidities for which the gradient boosting model outperformed the reference model by the largest margin (HF, CVD and CKD), the individuals predicted to be at highest risk by the gradient boosting model exhibited lower all-cause mortality incidence compared to their counterparts selected by the reference model (see Fig. 6b for the results for CKD and Supplementary Fig. 4 for the other four comorbidities). These results indicate that at least part of the performance advantage of the gradient boosting model over the reference model stems from its implicit ability to account for death risk in its predictions.

**Table 4.**
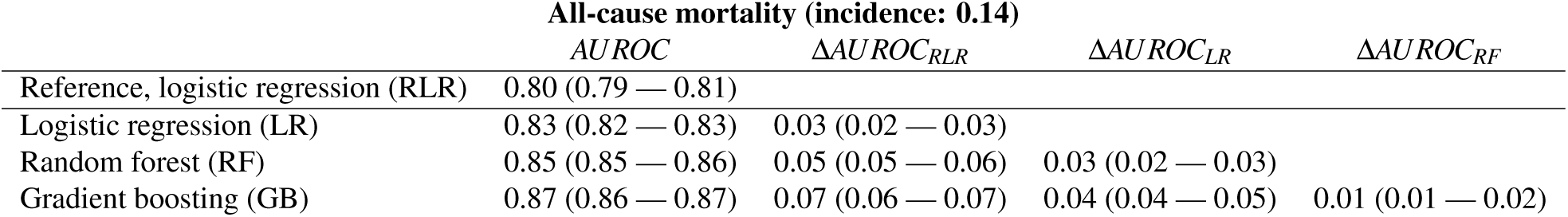
Area under receiver operating characteristic curve for prediction of all-cause mortality by the best reference and register-based models. Compared to the prediction of T2D comorbidities, all models achieved relatively high AUROCs, with all register-based models outperforming the reference model and gradient boosting outperforming the other register-based models. T2D, type 2 diabetes; AUROC, area under receiver operating characteristic curve.

**Figure 6.**
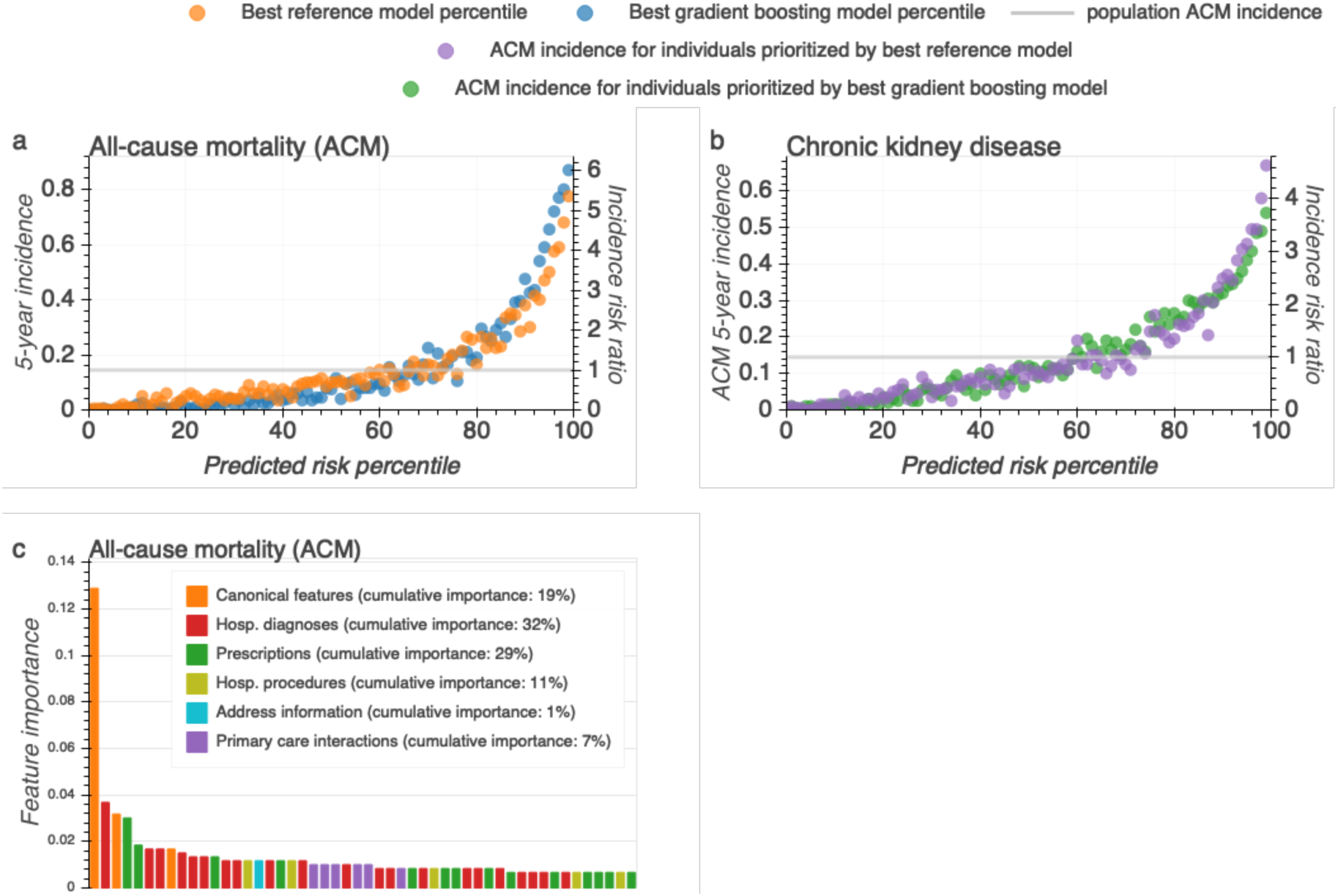
Death is a competing risk for diagnosing T2D comorbidities and its effect may be estimated by ML models implicitly. We investigated whether health register data and machine learning model could predict five-year risk of all-cause mortality (ACM) using the same procedure as for T2D comorbidities. (**a**) five-year incidence of ACM for population percentiles ranked by ACM risk predicted by gradient boosting (blue) and reference (orange) models. At the 95th percentile, both models identified individuals ranking multiples above the overall population incidence (ACM risk ratio of 4.53, and 3.46, respectively). (**b**) Incidence of ACM among population percentiles according to predicted risk of chronic kidney disease. Individuals stratified by the register-based model had a similar or lower risk of ACM than those binned into top percentiles by the reference model. (**c**) 50 register features most predictive of ACM according to the gradient boosting models feature importances. Unlike the case of the investigated T2D comorbidities, hospital diagnoses features were the most important followed by prescriptions second and canonical features third.

## Discussion

Here we evaluated the extent to which application of commonly used ML methods on comprehensive Danish nationwide health register data could predict selected T2D comorbidities, namely heart failure (HF), myocardial infarction (MI), stroke (ST), cardiovascular disease (CVD) and chronic kidney disease (CKD). Using the AUROC performance measure, we compared register-based models – logistic ridge regression, random forest and gradient boosting applied to nationwide information on hospital diagnoses, hospital procedures, drug prescriptions and claims information from primary care contractors – to a reference logistic ridge regression model based on age, sex, date of first T2D diagnosis and their interactions. For all comorbidities except ST, the register-based models outperformed the reference model. The gradient boosting model yielded the highest AUROCs significantly outperforming register-based logistic ridge regression for HF, CVD, and CKD as well as outperforming random forest models for all outcomes. For all register-based models, feature importances formed a long-tailed right-skewed distribution with the most important feature types being prescriptions, followed by the canonical features and hospital diagnoses. These results suggest that the predictive advantage of the register-based gradient boosting model stems from both the extensive inclusion of predictive features from health registers, especially drug prescriptions, as well as increased flexibility (capacity to learn complex, non-linear patterns in data) of the gradient boosting model over logistic ridge regression models.

The predictive advantage of the register-based gradient boosting models over the register-based logistic ridge regression stands in contrast to recent findings suggesting an absence of significant improvements from using ML techniques over logistic regression in clinical prediction tasks^13^. We believe that this improvement, as well as applicability of ML approaches on electronic health record data generally, depend on at least three factors. First, on the number of predictive features – our study used 6,181 features while the median number of features for studies presented by Christodoulou *et al.* was 19. Second, the presence of non-linear predictive patterns within the data favoring random forest and gradient boosting over logistic regression models. For example, in our study, the register-based gradient boosting did not outperform logistic ridge regression only for those outcomes (MI and ST) for which general trends, as explained by canonical predictive features (age, sex, date of diagnosis), were the most predictive (cumulative importance of 26% and 27%, respectively). Lastly, the number of samples sufficient for an ML model to learn the aforementioned complex patterns (the number of individuals in our study ranges from 175,460 to 200,646 as compared to 1,250 samples for the studies presented by Christodoulou *et al.*). Overall, these findings demonstrate that employment of ML techniques should only follow an earnest effort in application of simpler modelling techniques, such as logistic regression, as the potential performance gains from ML approaches may not outweigh the associated loss of interpretability or increased risk of over-fitting.

We note that the predictive performance of all models was relatively modest. The best AUROC performances were observed for gradient boosting for HF achieving an AUROC of 0.80 and CKD with an AUROC of 0.77, in both cases outperforming the reference model by DAUROC=0.05. Overall, the reference model achieved AUROCs from 0.62 for CVD to 0.75 for HF (95% CI) while gradient boosting achieved AUROCs from 0.69 for MI up to 0.81 for HF (95% CI). Focusing on the individuals predicted to be at highest risk, the register-based models were able to identify patients whose five-year T2D comorbidity risk was considerably increased, with risk ratios exceeding 3.1 for CVD, 3.8 for ST, 4.2 for MI, 4.8 for CKD and 5.5 for HF, which all were higher than the corresponding values for the reference model. Overall these results suggest that ML methods, such as gradient boosting, outperform or match logistic regression in prediction on nationwide health registers. Furthermore, with continued development and careful consideration, such an approach could be used to prioritize individuals for treatment regimens or to identify sub-groups of patients for whom an intervention could be cost-effective.

We found that AUROC values for prediction of comorbidities one-, two-, three- and four years after the first T2D diagnosis were comparable to prediction at the date of the first T2D diagnosis. This suggests that predictive register-derived risk factors for the investigated T2D comorbidities may already be present prior to given individual’s first T2D diagnosis. Similarly, while we observed a consistent increase in AUROC without using a time-split approach, these increases did not reach statistical significance, suggesting that changes in medical practice between 2000 and 2016 had only modest impact on the comorbidity diagnoses during that period. However, in our opinion, using a time-split approach is the more correct and principled way of evaluating prediction models in healthcare settings. For example, recent changes in medical practice, such as increased use of statins^65^ or introduction of new types of insulin or glucose-lowering drugs^66^ in the period of 2000 to 2013 could reduce the relevance of patterns learned from earlier predictions^44^ and, without proper evaluation, lead to overestimation of the models’ abilities to predict future events.

In this approach, individuals who died during the follow-up period either due to unrelated reasons or without an appropriate diagnosis in the death register were regarded as non-cases. Thus, death constituted a competing risk to the development of T2D comorbidities. However, due to their high flexibility, ML models could potentially account for this risk implicitly, i.e. by predicting individuals at higher risk of death prior to their diagnosis to be at lower risk of developing the T2D comorbidity. Indeed, in the analogous task of five-year all-cause mortality prediction at first T2D diagnosis, gradient boosting significantly outperformed the reference model achieving a relatively high AUROC of 0.87 when identifying subgroups at high (95th percentile risk ratio of 4.53) risk of death. Unlike their comorbidity counterparts, the best all-cause mortality prediction model was able to identify relatively rare but highly predictive hospital diagnosis features such as cancer diagnoses, suggesting that ML register-based approaches could be especially effective for all-cause mortality outcome prediction. Furthermore, for all models, all-cause mortality incidence was increasing alongside predicted comorbidity risk. However, for HF, CVD, and CKD individuals predicted to be at highest risk of developing a given comorbidity by a register-based model had a lower incidence of all-cause mortality than their counterparts selected by the reference model. This indicates that ML register-based models can outperform the reference model through better estimation of individuals’ risk of death and weighing it against the risk of a given comorbidity.

Employment of Danish population-wide health register data for prediction of health-related outcome has a number of advantages. Firstly, the registers cover virtually every Danish citizen, which reduces the risk of training models based on unequal representations of population sub-groups and minimizes censoring. Secondly, our analyses include information from multiple Danish registers spanning from hospitalization events to drug prescriptions and claims from primary care contractors. Thus, our results give a more realistic picture of the extent to which register data can aid in the prediction of health-related outcomes, in our case T2D comorbidities. The observation that the register-based models exhibited consistently higher performances compared to the reference models underlines that register-based models may represent powerful frameworks to identify individuals at high risk of developing T2D comorbidities. Taking into account eligibility criteria, our approach can be used to prioritize individuals for treatment regimens or to identify sub-groups of patients for whom an intervention could become more cost-effective. Furthermore, due to the nationwide breadth of this data, our models could be retrained with genetics data, socioeconomic information, lab measurements and/or other comprehensive molecular data types and electronic medical record systems to move a step closer towards being incorporated into the routine care of patients newly diagnosed with T2D. The proof-of-concept approach presented in this work could be well suited for a prediction of outcomes secondary to a wide array of diseases including cardiovascular and hematological diseases and potentially cognitive and psychiatric disorders.

Despite its strengths, our work has a number of important shortcomings. Firstly, due to the lack of diagnosis information from general practitioners, we rely on hospital diagnoses to identify the onset of investigated T2D comorbidities. In doing so, we possibly include as cases individuals who have already been diagnosed by their general practitioner and potentially were already receiving treatment at the time of prediction. Our models could then identify the treatment-related prescriptions as highly predictive of a future hospital diagnosis of the comorbidity. This may be especially true for CVD, for which we observed a prescription of cardiovascular system drugs as one of the most predictive features. Similarly, some of the individuals predicted to be at highest risk are already known to be in the high-risk group. For example, for CKD the best model also identified as highly predictive prescription of calcium blockers – agents typically prescribed to individuals with hypertension who because of this diagnosis are at risk of kidney disease. This indicates that some of the at-risk individuals were known to be at risk for CKD before their first T2D diagnosis. In future work, these confounders could be addressed by filtering out individuals with medical prescriptions indicative of the comorbidity at the time of prediction. Secondly, logistic ridge regression, random forest and gradient boosting are regarded as interpretable models – that is models which predictions can be analyzed to gain an understanding of their predictions. However, the highly correlated nature of register features makes direct interpretation of feature importance challenging. This deficiency could be partially addressed by approaches such as LIME^67^, ANCHORS^68^, SHAP^69^ or LRP^70^ which offer a promise of identifying the most predictive features for each individual separately as opposed to feature importance averaged across all individuals. Analysis of patterns in individual-level feature importance could reveal non-linear relationships between features as well as patient sub-groups with different risk factor profiles. Finally, we acknowledge that future work should include additional models, extended parametrization and use other frameworks^71^ and performance measures^72^.

Consequently, there is a number of additional aspects of the proposed framework that could be improved. Firstly, our approach did not leverage the temporal dimension in the data, potentially missing predictive information stored in order or time distribution of events. Recently, studies which applied models incorporating time to electronic health record data suggest that inclusion of temporal information improves prediction^21, 73^. Secondly, instead of providing binary prediction (i.e. predicting whether an individual will be diagnosed with an outcome or not within a certain time period) the model could perform probabilistic estimation of time to outcome diagnosis. By additionally estimating a date of an adverse event, models predictions could better inform the type or intensity of an intervention. Similarly, future models could additionally estimate time and risk of death which could inform intervention considerations and simplify the learning task. Lastly, the proposed framework could be easily extended to leverage additional sources of information such as molecular data, lab measurements and socioeconomic information to improve its prediction.

In summary, we show that inclusion of longitudinal Danish register data comprising drug prescriptions, hospital diagnoses, hospital procedures and claims from primary care contractors moderately improves prediction of T2D comorbidities compared to a reference model based on readily-available canonical predictors. Our findings indicate that Danish registers may be used to develop nationwide *in silico* screening approaches to identify chronic disease patients at risk of developing a certain comorbidity, but stress that, most likely, additional data – for instance genotyping, lab measurements or socio-economic data – is needed to more robustly accomplish that goal.

## Acknowledgments

This work was supported by a research grant from the Danish Diabetes Academy funded by the Novo Nordisk Foundation. Furthermore, THP and PD acknowledge the Novo Nordisk Foundation (grant number NNF18CC0034900) and Lundbeck Foundation (grant number R190-2014-3904 to THP). We thank Jonatan Thompson, Andreas Rieckmann and Asker Brejnrod for their comments, conducive discussion and insights.

## Additional information

### Ethical approval

The project was approved by Danish Data Protection Agency (2015-57-0102). Under Danish law register-based research does not require informed consent.

### Data availability

The Danish health register data used in this study is available for research through Statistics Denmark^74^ and the Danish Health Data Authority^75^.

### Contributions

PD and THP conceived this study. PD performed all the analyses. TAG, KR and HH provided methodological suggestions. PD, THP and MA wrote the manuscript. All authors reviewed and commented on the manuscript.

