## Supplementary Material for "Nationwide prediction of type 2 diabetes comorbidities"

for

**This file includes:**

- Supplementary Figures 1 to 4
- Supplementary Tables 1 to 7

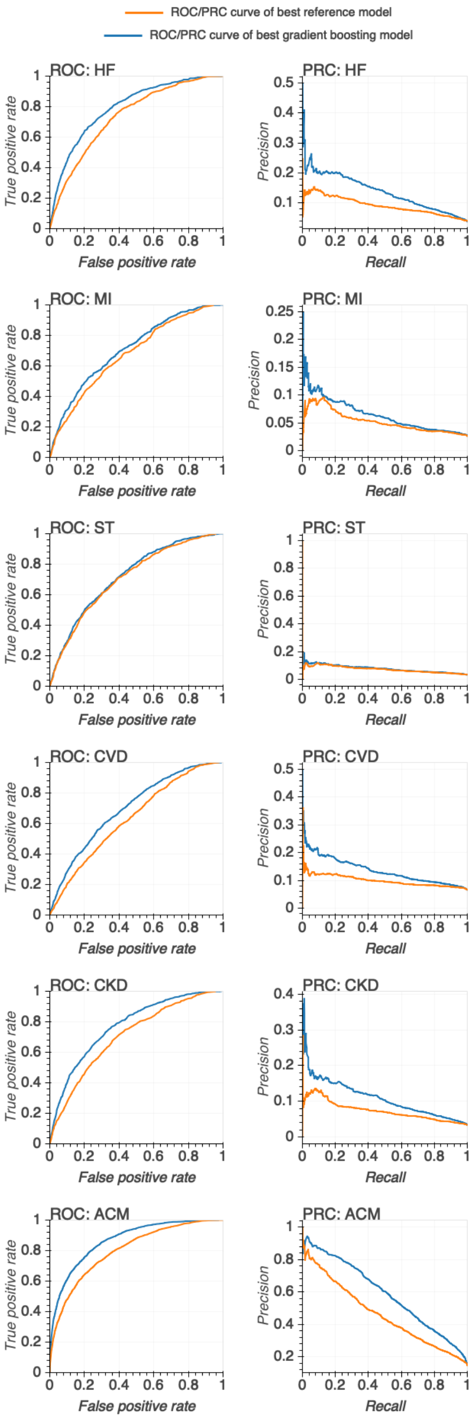


**Supplementary Figure 1.** Receiver operating characteristic (ROC) and precision-recall (PRC) curves for best parametrization of baseline (orange) and gradient boosting (blue) models. HF, heart failure; MI, myocardial infarction; ST, stroke; CVD, cardiovascular disease; CKD, chronic kidney disease; ACM, all-cause mortality.

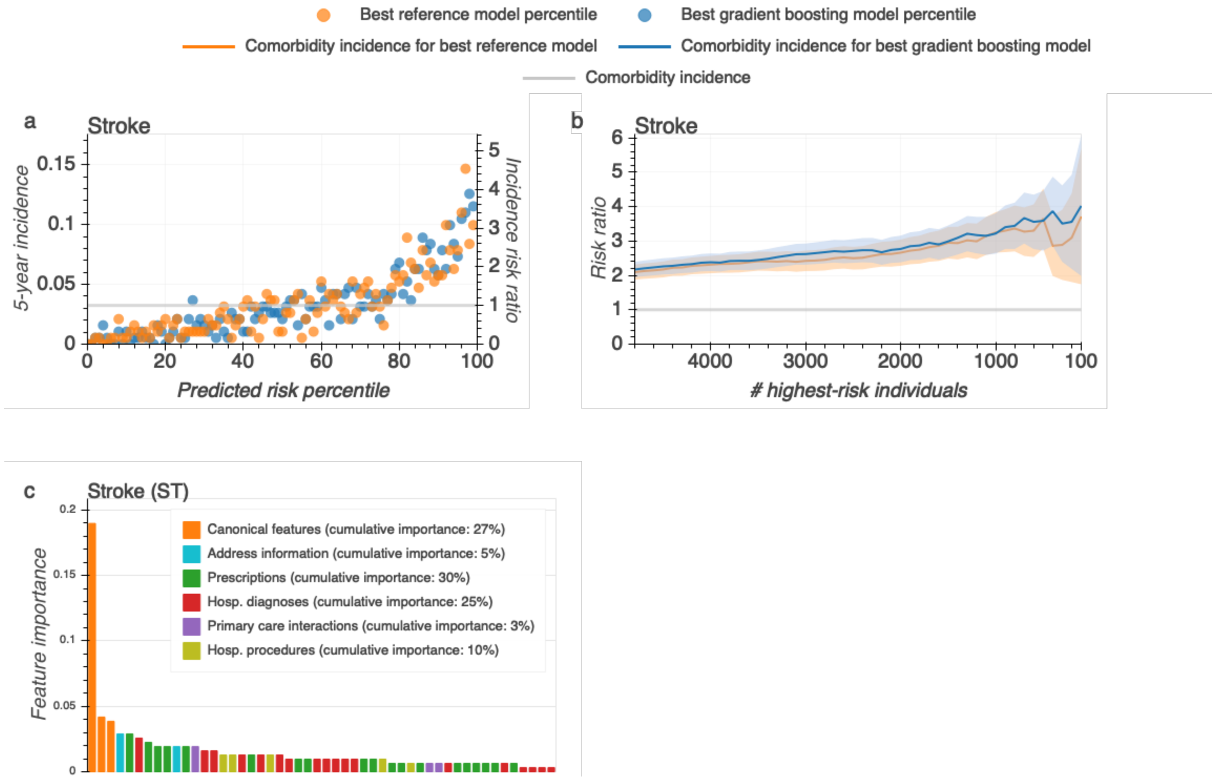


**Supplementary Figure 2.** (**a**) five-year incidence of hospital diagnosis of stroke for population percentiles ranked by risk as predicted by the best gradient boosting (blue) and the best baseline (orange) models. (**b**) Individuals were ranked according to their predicted risk of stroke by the best gradient boosting (blue) and the best baseline (orange) models. For a number of thresholds, shown are risk ratios, calculated as the stroke incidence of individuals ranking above that thresholds over ST incidence in the entire study population. 95% confidence interval (shaded areas) were obtained through bootstrap sampling. (**c**) 50 most predictive features for stroke according to the best gradient boosting models feature importances.

| | **T2D population** | **HF** | **MI** |
|---|---|---|---|
| # Individuals | 181,100 | 169,729 | 169,972 |
| # Cases | - | 8,222 (4.84%) | 6,039 (3.55%) |
| # Non-cases | - | 161,507 (95.16%) | 163,933 (96.45%) |
| % Women | 46.98 | 47.53 | 48.13 |
| Median age at T2D diagnosis | 61.71 | 60.95 | 61.24 |
| # Days until outcome | - | 1672.90 | 1546.60 |
| # Features in RFV | - | 6,155 | 6,155 |
| | **ST** | **CVD** | **CKD** |
| # Individuals | 169,870 | 154,909 | 178,586 |
| # Cases | 7,467 (4.40%) | 11,880 (7.67%) | 5,361 (3.00%) |
| # Non-cases | 162,403 (95.60%) | 143,029 (92.33%) | 173,225 (97.00%) |
| % Women | 47.37 | 49.11 | 47.13 |
| Median age at T2D diagnosis | 61.07 | 60.41 | 61.61 |
| # Days until outcome | 1636.50 | 1513.50 | 2060.20 |
| # Features in RFV | 6,155 | 6,155 | 6,155 |

**(a)** one year after first T2D diagnosis

| | **T2D population** | **HF** | **MI** |
|---|---|---|---|
| # Individuals | 165,482 | 154,684 | 154,997 |
| # Cases | - | 8,019 (5.18%) | 5,814 (3.75%) |
| # Non-cases | - | 146,665 (94.82%) | 149,183 (96.25%) |
| % Women | 46.89 | 47.42 | 48.05 |
| Median age at T2D diagnosis | 62.35 | 61.56 | 61.85 |
| # Days until outcome | - | 1659.20 | 1562.80 |
| # Features in RFV | - | 6,155 | 6,155 |
| | **ST** | **CVD** | **CKD** |
| # Individuals | 154,683 | 140,467 | 163,069 |
| # Cases | 7,232 (4.68%) | 11,156 (7.94%) | 5,318 (3.26%) |
| # Non-cases | 147,451 (95.32%) | 129,311 (92.06%) | 157,751 (96.74%) |
| % Women | 47.28 | 49.05 | 47.04 |
| Median age at T2D diagnosis | 61.66 | 61.04 | 62.25 |
| # Days until outcome | 1625.90 | 1503.60 | 2036.50 |
| # Features in RFV | 6,155 | 6,155 | 6,155 |

**(b)** two years after first T2D diagnosis

| | **T2D population** | **HF** | **MI** |
|---|---|---|---|
| # Individuals | 150,228 | 140,053 | 140,418 |
| # Cases | - | 7,667 (5.47%) | 5,492 (3.91%) |
| # Non-cases | - | 132,386 (94.53%) | 134,926 (96.09%) |
| % Women | 46.73 | 47.26 | 47.88 |
| Median age at T2D diagnosis | 62.90 | 62.11 | 62.41 |
| # Days until outcome | - | 1652.20 | 1560.40 |
| # Features in RFV | - | 6,155 | 6,155 |
| | **ST** | **CVD** | **CKD** |
| # Individuals | 139,877 | 126,650 | 147,830 |
| # Cases | 6,896 (4.93%) | 10,452 (8.25%) | 5,178 (3.50%) |
| # Non-cases | 132,981 (95.07%) | 116,198 (91.75%) | 142,652 (96.50%) |
| % Women | 47.18 | 48.84 | 46.91 |
| Median age at T2D diagnosis | 62.22 | 61.60 | 62.78 |
| # Days until outcome | 1592.60 | 1494.80 | 2012.60 |
| # Features in RFV | 6,155 | 6,155 | 6,155 |

**(c)** three years after first T2D diagnosis

| | **T2D population** | **HF** | **MI** |
|---|---|---|---|
| # Individuals | 137,240 | 127,580 | 127,930 |
| # Cases | - | 7,403 (5.80%) | 5,298 (4.14%) |
| # Non-cases | - | 120,177 (94.20%) | 122,632 (95.86%) |
| % Women | 46.38 | 46.89 | 47.51 |
| Median age at T2D diagnosis | 63.56 | 62.75 | 63.07 |
| # Days until outcome | - | 1625.90 | 1543.10 |
| # Features in RFV | - | 6,155 | 6,155 |
| | **ST** | **CVD** | **CKD** |
| # Individuals | 127,240 | 114,754 | 134,874 |
| # Cases | 6,666 (5.24%) | 9,926 (8.65%) | 5,070 (3.76%) |
| # Non-cases | 120,574 (94.76%) | 104,828 (91.35%) | 129,804 (96.24%) |
| % Women | 46.83 | 48.47 | 46.55 |
| Median age at T2D diagnosis | 62.84 | 62.26 | 63.44 |
| # Days until outcome | 1571.40 | 1474.40 | 2018.10 |
| # Features in RFV | 6,155 | 6,155 | 6,155 |

**(d)** four years after first T2D diagnosis

**Supplementary Table 1.** Population characteristics at different dates of prediction. T2D, type 2 diabetes; HF, heart failure; MI, myocardial infarction; ST, stroke; CVD, cardiovascular disease; CKD, chronic kidney disease; RFV, register feature vector; #, number of.

**Heart failure**

| Model | Hyperparameters | AUROC |
|---|---|---|
| GB | max_depth: 2 learning_rate: 0.1 n_estimators: 200 | 0.803 |
| GB | max_depth: 3 learning_rate: 0.05 n_estimators: 200 | 0.803 |
| GB | max_depth: 4 learning_rate: 0.05 n_estimators: 200 | 0.803 |
| GB | max_depth: 3 learning_rate: 0.1 n_estimators: 200 | 0.803 |
| GB | max_depth: 2 learning_rate: 0.05 n_estimators: 200 | 0.802 |
| GB | max_depth: 4 learning_rate: 0.025 n_estimators: 200 | 0.802 |
| GB | max_depth: 4 learning_rate: 0.1 n_estimators: 200 | 0.802 |
| GB | max_depth: 5 learning_rate: 0.025 n_estimators: 200 | 0.801 |
| GB | max_depth: 5 learning_rate: 0.05 n_estimators: 200 | 0.801 |
| GB | max_depth: 3 learning_rate: 0.025 n_estimators: 200 | 0.801 |
| GB | max_depth: 5 learning_rate: 0.1 n_estimators: 200 | 0.800 |
| GB | max_depth: 2 learning_rate: 0.025 n_estimators: 200 | 0.798 |
| RF | n_estimators: 1500 max_depth: 12 | 0.790 |
| RF | n_estimators: 1000 max_depth: 12 | 0.790 |
| RF | n_estimators: 1500 max_depth: 14 | 0.787 |
| RF | n_estimators: 1000 max_depth: 14 | 0.786 |
| LR | C: 0.8 max_iter: 500 | 0.782 |
| LR | C: 0.6 max_iter: 500 | 0.782 |
| LR | C: 0.7 max_iter: 500 | 0.781 |
| RF | n_estimators: 1500 max_depth: 16 | 0.781 |
| RF | n_estimators: 1000 max_depth: 16 | 0.781 |
| LR | C: 0.7 max_iter: 400 | 0.780 |
| LR | C: 0.6 max_iter: 400 | 0.780 |
| LR | C: 0.8 max_iter: 400 | 0.780 |
| LR | C: 0.6 max_iter: 300 | 0.779 |
| LR | C: 0.7 max_iter: 300 | 0.778 |
| LR | C: 0.8 max_iter: 300 | 0.778 |
| RLR | C: 0.6 max_iter: 300 | 0.748 |
| RLR | C: 0.6 max_iter: 400 | 0.748 |
| RLR | C: 0.6 max_iter: 500 | 0.748 |
| RLR | C: 0.7 max_iter: 300 | 0.748 |
| RLR | C: 0.7 max_iter: 400 | 0.748 |
| RLR | C: 0.7 max_iter: 500 | 0.748 |
| RLR | C: 0.8 max_iter: 300 | 0.748 |
| RLR | C: 0.8 max_iter: 400 | 0.748 |
| RLR | C: 0.8 max_iter: 500 | 0.748 |

**Myocardial infarction**

| Model | Hyperparameters | AUROC |
|---|---|---|
| GB | max_depth: 2 learning_rate: 0.1 n_estimators: 200 | 0.730 |
| GB | max_depth: 2 learning_rate: 0.05 n_estimators: 200 | 0.730 |
| GB | max_depth: 3 learning_rate: 0.025 n_estimators: 200 | 0.729 |
| GB | max_depth: 3 learning_rate: 0.1 n_estimators: 200 | 0.729 |
| GB | max_depth: 4 learning_rate: 0.025 n_estimators: 200 | 0.729 |
| GB | max_depth: 3 learning_rate: 0.05 n_estimators: 200 | 0.729 |
| GB | max_depth: 4 learning_rate: 0.05 n_estimators: 200 | 0.728 |
| GB | max_depth: 4 learning_rate: 0.1 n_estimators: 200 | 0.727 |
| GB | max_depth: 2 learning_rate: 0.025 n_estimators: 200 | 0.727 |
| GB | max_depth: 5 learning_rate: 0.025 n_estimators: 200 | 0.727 |
| GB | max_depth: 5 learning_rate: 0.05 n_estimators: 200 | 0.726 |
| GB | max_depth: 5 learning_rate: 0.1 n_estimators: 200 | 0.725 |
| LR | C: 0.6 max_iter: 400 | 0.723 |
| LR | C: 0.8 max_iter: 500 | 0.722 |
| LR | C: 0.6 max_iter: 500 | 0.722 |
| LR | C: 0.7 max_iter: 500 | 0.722 |
| LR | C: 0.7 max_iter: 400 | 0.722 |
| LR | C: 0.8 max_iter: 300 | 0.722 |
| LR | C: 0.8 max_iter: 400 | 0.722 |
| LR | C: 0.6 max_iter: 300 | 0.721 |
| LR | C: 0.7 max_iter: 300 | 0.721 |
| RF | n_estimators: 1500 max_depth: 12 | 0.712 |
| RF | n_estimators: 1000 max_depth: 12 | 0.712 |
| RLR | C: 0.6 max_iter: 300 | 0.703 |
| RLR | C: 0.6 max_iter: 400 | 0.703 |
| RLR | C: 0.6 max_iter: 500 | 0.703 |
| RLR | C: 0.7 max_iter: 300 | 0.703 |
| RLR | C: 0.7 max_iter: 400 | 0.703 |
| RLR | C: 0.7 max_iter: 500 | 0.703 |
| RLR | C: 0.8 max_iter: 300 | 0.703 |
| RLR | C: 0.8 max_iter: 400 | 0.703 |
| RLR | C: 0.8 max_iter: 500 | 0.703 |
| RF | n_estimators: 1000 max_depth: 14 | 0.703 |
| RF | n_estimators: 1500 max_depth: 14 | 0.703 |
| RF | n_estimators: 1000 max_depth: 16 | 0.691 |
| RF | n_estimators: 1500 max_depth: 16 | 0.691 |

**Stroke**

| Model | Hyperparameters | AUROC |
|---|---|---|
| GB | max_depth: 2 learning_rate: 0.1 n_estimators: 200 | 0.734 |
| GB | max_depth: 2 learning_rate: 0.05 n_estimators: 200 | 0.733 |
| GB | max_depth: 3 learning_rate: 0.05 n_estimators: 200 | 0.733 |
| GB | max_depth: 3 learning_rate: 0.1 n_estimators: 200 | 0.733 |
| GB | max_depth: 3 learning_rate: 0.025 n_estimators: 200 | 0.733 |
| GB | max_depth: 4 learning_rate: 0.025 n_estimators: 200 | 0.732 |
| GB | max_depth: 4 learning_rate: 0.05 n_estimators: 200 | 0.731 |
| GB | max_depth: 4 learning_rate: 0.1 n_estimators: 200 | 0.731 |
| GB | max_depth: 2 learning_rate: 0.025 n_estimators: 200 | 0.730 |
| GB | max_depth: 5 learning_rate: 0.025 n_estimators: 200 | 0.730 |
| GB | max_depth: 5 learning_rate: 0.05 n_estimators: 200 | 0.729 |
| GB | max_depth: 5 learning_rate: 0.1 n_estimators: 200 | 0.728 |
| LR | C: 0.8 max_iter: 500 | 0.726 |
| LR | C: 0.6 max_iter: 300 | 0.725 |
| LR | C: 0.8 max_iter: 400 | 0.725 |
| LR | C: 0.6 max_iter: 500 | 0.725 |
| LR | C: 0.6 max_iter: 400 | 0.725 |
| LR | C: 0.8 max_iter: 300 | 0.725 |
| LR | C: 0.7 max_iter: 300 | 0.725 |
| LR | C: 0.7 max_iter: 500 | 0.725 |
| LR | C: 0.7 max_iter: 400 | 0.725 |
| RLR | C: 0.6 max_iter: 300 | 0.717 |
| RLR | C: 0.6 max_iter: 400 | 0.717 |
| RLR | C: 0.6 max_iter: 500 | 0.717 |
| RLR | C: 0.7 max_iter: 300 | 0.717 |
| RLR | C: 0.7 max_iter: 400 | 0.717 |
| RLR | C: 0.7 max_iter: 500 | 0.717 |
| RLR | C: 0.8 max_iter: 300 | 0.717 |
| RLR | C: 0.8 max_iter: 400 | 0.717 |
| RLR | C: 0.8 max_iter: 500 | 0.717 |
| RF | n_estimators: 1500 max_depth: 12 | 0.713 |
| RF | n_estimators: 1000 max_depth: 12 | 0.713 |
| RF | n_estimators: 1500 max_depth: 14 | 0.707 |
| RF | n_estimators: 1000 max_depth: 14 | 0.707 |
| RF | n_estimators: 1500 max_depth: 16 | 0.699 |
| RF | n_estimators: 1000 max_depth: 16 | 0.699 |

**Cardiovascular disease**

| Model | Hyperparameters | AUROC |
|---|---|---|
| GB | max_depth: 2 learning_rate: 0.1 n_estimators: 200 | 0.726 |
| GB | max_depth: 3 learning_rate: 0.05 n_estimators: 200 | 0.726 |
| GB | max_depth: 4 learning_rate: 0.05 n_estimators: 200 | 0.726 |
| GB | max_depth: 3 learning_rate: 0.1 n_estimators: 200 | 0.726 |
| GB | max_depth: 4 learning_rate: 0.025 n_estimators: 200 | 0.724 |
| GB | max_depth: 4 learning_rate: 0.1 n_estimators: 200 | 0.724 |
| GB | max_depth: 5 learning_rate: 0.05 n_estimators: 200 | 0.724 |
| GB | max_depth: 2 learning_rate: 0.05 n_estimators: 200 | 0.724 |
| GB | max_depth: 5 learning_rate: 0.025 n_estimators: 200 | 0.723 |
| GB | max_depth: 5 learning_rate: 0.1 n_estimators: 200 | 0.723 |
| GB | max_depth: 3 learning_rate: 0.025 n_estimators: 200 | 0.722 |
| RF | n_estimators: 1500 max_depth: 12 | 0.707 |
| RF | n_estimators: 1000 max_depth: 12 | 0.707 |
| RF | n_estimators: 1500 max_depth: 14 | 0.703 |
| RF | n_estimators: 1000 max_depth: 14 | 0.703 |
| GB | max_depth: 2 learning_rate: 0.025 n_estimators: 200 | 0.703 |
| RF | n_estimators: 1500 max_depth: 16 | 0.696 |
| RF | n_estimators: 1000 max_depth: 16 | 0.696 |
| LR | C: 0.7 max_iter: 500 | 0.691 |
| LR | C: 0.7 max_iter: 400 | 0.690 |
| LR | C: 0.6 max_iter: 500 | 0.690 |
| LR | C: 0.8 max_iter: 500 | 0.689 |
| LR | C: 0.7 max_iter: 300 | 0.688 |
| LR | C: 0.8 max_iter: 400 | 0.688 |
| LR | C: 0.6 max_iter: 300 | 0.688 |
| LR | C: 0.6 max_iter: 400 | 0.688 |
| LR | C: 0.8 max_iter: 300 | 0.688 |
| RLR | C: 0.6 max_iter: 300 | 0.665 |
| RLR | C: 0.6 max_iter: 400 | 0.665 |
| RLR | C: 0.6 max_iter: 500 | 0.665 |
| RLR | C: 0.7 max_iter: 300 | 0.665 |
| RLR | C: 0.7 max_iter: 400 | 0.665 |
| RLR | C: 0.7 max_iter: 500 | 0.665 |
| RLR | C: 0.8 max_iter: 300 | 0.665 |
| RLR | C: 0.8 max_iter: 400 | 0.665 |
| RLR | C: 0.8 max_iter: 500 | 0.665 |

**Supplementary Table 2.** Area under receiver operating characteristic curve (AUROC) performance for each tested parametrization for heart failure, myocardial infarction, stroke and cardiovascular disease. Date of prediction was set to date of first type 2 diabetes diagnosis. Hyperparameter names correspond to parameter names from the python sklearn package. GB, gradient boosting; RF, random forest; LR, register-based logistic regression; RLR, reference logistic regression.

**Chronic kidney disease**

| Model | Hyperparameters | AUROC |
|---|---|---|
| GB | max_depth: 2 learning_rate: 0.1 n_estimators: 200 | 0.754 |
| GB | max_depth: 3 learning_rate: 0.05 n_estimators: 200 | 0.754 |
| GB | max_depth: 2 learning_rate: 0.05 n_estimators: 200 | 0.753 |
| GB | max_depth: 3 learning_rate: 0.1 n_estimators: 200 | 0.753 |
| GB | max_depth: 3 learning_rate: 0.025 n_estimators: 200 | 0.752 |
| GB | max_depth: 4 learning_rate: 0.05 n_estimators: 200 | 0.752 |
| GB | max_depth: 4 learning_rate: 0.025 n_estimators: 200 | 0.751 |
| GB | max_depth: 2 learning_rate: 0.025 n_estimators: 200 | 0.749 |
| GB | max_depth: 4 learning_rate: 0.1 n_estimators: 200 | 0.749 |
| GB | max_depth: 5 learning_rate: 0.025 n_estimators: 200 | 0.749 |
| GB | max_depth: 5 learning_rate: 0.05 n_estimators: 200 | 0.747 |
| GB | max_depth: 5 learning_rate: 0.1 n_estimators: 200 | 0.745 |
| LR | C: 0.6 max_iter: 500 | 0.730 |
| LR | C: 0.6 max_iter: 400 | 0.729 |
| RF | n_estimators: 1500 max_depth: 12 | 0.729 |
| LR | C: 0.8 max_iter: 400 | 0.729 |
| LR | C: 0.8 max_iter: 500 | 0.729 |
| RF | n_estimators: 1000 max_depth: 12 | 0.729 |
| LR | C: 0.7 max_iter: 500 | 0.727 |
| LR | C: 0.7 max_iter: 400 | 0.727 |
| LR | C: 0.6 max_iter: 300 | 0.726 |
| LR | C: 0.7 max_iter: 300 | 0.726 |
| LR | C: 0.8 max_iter: 300 | 0.724 |
| RF | n_estimators: 1500 max_depth: 14 | 0.718 |
| RF | n_estimators: 1000 max_depth: 14 | 0.717 |
| RF | n_estimators: 1500 max_depth: 16 | 0.701 |
| RF | n_estimators: 1000 max_depth: 16 | 0.700 |
| RLR | C: 0.6 max_iter: 300 | 0.693 |
| RLR | C: 0.6 max_iter: 400 | 0.693 |
| RLR | C: 0.6 max_iter: 500 | 0.693 |
| RLR | C: 0.7 max_iter: 300 | 0.693 |
| RLR | C: 0.7 max_iter: 400 | 0.693 |
| RLR | C: 0.7 max_iter: 500 | 0.693 |
| RLR | C: 0.8 max_iter: 300 | 0.693 |
| RLR | C: 0.8 max_iter: 400 | 0.693 |
| RLR | C: 0.8 max_iter: 500 | 0.693 |

**All-cause mortality**

| Model | Hyperparameters | AUROC |
|---|---|---|
| GB | max_depth: 4 learning_rate: 0.1 n_estimators: 200 | 0.864 |
| GB | max_depth: 5 learning_rate: 0.1 n_estimators: 200 | 0.863 |
| GB | max_depth: 3 learning_rate: 0.1 n_estimators: 200 | 0.863 |
| GB | max_depth: 5 learning_rate: 0.05 n_estimators: 200 | 0.863 |
| GB | max_depth: 4 learning_rate: 0.05 n_estimators: 200 | 0.862 |
| GB | max_depth: 2 learning_rate: 0.1 n_estimators: 200 | 0.861 |
| GB | max_depth: 3 learning_rate: 0.05 n_estimators: 200 | 0.860 |
| GB | max_depth: 5 learning_rate: 0.025 n_estimators: 200 | 0.859 |
| GB | max_depth: 4 learning_rate: 0.025 n_estimators: 200 | 0.857 |
| GB | max_depth: 2 learning_rate: 0.05 n_estimators: 200 | 0.856 |
| GB | max_depth: 3 learning_rate: 0.025 n_estimators: 200 | 0.854 |
| GB | max_depth: 2 learning_rate: 0.025 n_estimators: 200 | 0.847 |
| RF | n_estimators: 1500 max_depth: 16 | 0.845 |
| RF | n_estimators: 1000 max_depth: 16 | 0.845 |
| RF | n_estimators: 1500 max_depth: 14 | 0.842 |
| RF | n_estimators: 1000 max_depth: 14 | 0.842 |
| RF | n_estimators: 1500 max_depth: 12 | 0.838 |
| RF | n_estimators: 1000 max_depth: 12 | 0.838 |
| LR | C: 0.8 max_iter: 500 | 0.827 |
| LR | C: 0.7 max_iter: 500 | 0.827 |
| LR | C: 0.6 max_iter: 500 | 0.826 |
| LR | C: 0.8 max_iter: 400 | 0.826 |
| LR | C: 0.7 max_iter: 400 | 0.826 |
| LR | C: 0.6 max_iter: 400 | 0.825 |
| LR | C: 0.8 max_iter: 300 | 0.823 |
| LR | C: 0.6 max_iter: 300 | 0.822 |
| LR | C: 0.7 max_iter: 300 | 0.822 |
| RLR | C: 0.6 max_iter: 300 | 0.796 |
| RLR | C: 0.6 max_iter: 400 | 0.796 |
| RLR | C: 0.6 max_iter: 500 | 0.796 |
| RLR | C: 0.7 max_iter: 300 | 0.796 |
| RLR | C: 0.7 max_iter: 400 | 0.796 |
| RLR | C: 0.7 max_iter: 500 | 0.796 |
| RLR | C: 0.8 max_iter: 300 | 0.796 |
| RLR | C: 0.8 max_iter: 400 | 0.796 |
| RLR | C: 0.8 max_iter: 500 | 0.796 |

**Supplementary Table 3.** Area under receiver operating characteristic curve (AUROC) performance for each tested parametrization for chronic kidney disease and all cause mortality. Date of prediction was set to date of first type 2 diabetes diagnosis. Hyperparameter names correspond to parameter names from the the python sklearn package. GB, gradient boosting; RF, random forest; RLR, register-based logistic regression; LR, reference logistic regression.

**Heart failure (incidence: 0.04)**

| | AUROC |
|---|---|
| Reference, logistic regression | 0.74 (0.72 — 0.76) |
| Gradient boosting | 0.80 (0.78 — 0.81) |

**Myocardial infarction (incidence: 0.03)**

| | AUROC |
|---|---|
| Reference, logistic regression | 0.68 (0.66 — 0.71) |
| Gradient boosting | 0.73 (0.71 — 0.75) |

**Stroke (incidence: 0.03)**

| | AUROC |
|---|---|
| Reference, logistic regression | 0.70 (0.68 — 0.72) |
| Gradient boosting | 0.75 (0.73 — 0.77) |

**Cardiovascular disease (incidence: 0.06)**

| | AUROC |
|---|---|
| Reference, logistic regression | 0.64 (0.62 — 0.65) |
| Gradient boosting | 0.71 (0.70 — 0.73) |

**Chronic kidney disease (incidence: 0.03)**

| | AUROC |
|---|---|
| Reference, logistic regression | 0.68 (0.66 — 0.70) |
| Gradient boosting | 0.77 (0.75 — 0.78) |

**(a)** one years after first T2D diagnosis

**Heart failure (incidence: 0.04)**

| | AUROC |
|---|---|
| Reference, logistic regression | 0.74 (0.72 — 0.76) |
| Gradient boosting | 0.80 (0.78 — 0.81) |

**Myocardial infarction (incidence: 0.03)**

| | AUROC |
|---|---|
| Reference, logistic regression | 0.69 (0.67 — 0.71) |
| Gradient boosting | 0.74 (0.71 — 0.76) |

**Stroke (incidence: 0.04)**

| | AUROC |
|---|---|
| Reference, logistic regression | 0.71 (0.69 — 0.73) |
| Gradient boosting | 0.75 (0.73 — 0.77) |

**Cardiovascular disease (incidence: 0.06)**

| | AUROC |
|---|---|
| Reference, logistic regression | 0.65 (0.64 — 0.67) |
| Gradient boosting | 0.72 (0.71 — 0.74) |

**Chronic kidney disease (incidence: 0.04)**

| | AUROC |
|---|---|
| Reference, logistic regression | 0.70 (0.68 — 0.72) |
| Gradient boosting | 0.77 (0.75 — 0.79) |

**(b)** two years after first T2D diagnosis

**Heart failure (incidence: 0.04)**

| | AUROC |
|---|---|
| Reference, logistic regression | 0.75 (0.73 — 0.77) |
| Gradient boosting | 0.81 (0.79 — 0.83) |

**Myocardial infarction (incidence: 0.03)**

| | AUROC |
|---|---|
| Reference, logistic regression | 0.69 (0.67 — 0.72) |
| Gradient boosting | 0.72 (0.70 — 0.75) |

**Stroke (incidence: 0.04)**

| | AUROC |
|---|---|
| Reference, logistic regression | 0.72 (0.70 — 0.74) |
| Gradient boosting | 0.75 (0.73 — 0.77) |

**Cardiovascular disease (incidence: 0.06)**

| | AUROC |
|---|---|
| Reference, logistic regression | 0.65 (0.63 — 0.67) |
| Gradient boosting | 0.72 (0.70 — 0.73) |

**Chronic kidney disease (incidence: 0.04)**

| | AUROC |
|---|---|
| Reference, logistic regression | 0.72 (0.70 — 0.74) |
| Gradient boosting | 0.79 (0.77 — 0.80) |

**(c)** three years after first T2D diagnosis

**Heart failure (incidence: 0.04)**

| | AUROC |
|---|---|
| Reference, logistic regression | 0.74 (0.72 — 0.75) |
| Gradient boosting | 0.81 (0.79 — 0.82) |

**Myocardial infarction (incidence: 0.03)**

| | AUROC |
|---|---|
| Reference, logistic regression | 0.68 (0.66 — 0.71) |
| Gradient boosting | 0.73 (0.70 — 0.75) |

**Stroke (incidence: 0.04)**

| | AUROC |
|---|---|
| Reference, logistic regression | 0.72 (0.70 — 0.74) |
| Gradient boosting | 0.75 (0.73 — 0.77) |

**Cardiovascular disease (incidence: 0.06)**

| | AUROC |
|---|---|
| Reference, logistic regression | 0.64 (0.62 — 0.66) |
| Gradient boosting | 0.71 (0.69 — 0.73) |

**Chronic kidney disease (incidence: 0.04)**

| | AUROC |
|---|---|
| Reference, logistic regression | 0.70 (0.68 — 0.72) |
| Gradient boosting | 0.79 (0.77 — 0.81) |

**(d)** four years after first T2D diagnosis

**Supplementary Table 4.** Gradient boosting and baseline model prediction performance for different dates of prediction. Incidence is the proportion of cases within the population. AUROC is area under receiver operating characteristic curve which confidence intervals were obtained by bootstrap sampling procedure. T2D, type 2 diabetes.

**Heart failure**

| $t_P$ | $I$ | Gradient Boosting AUROC | Reference AUROC |
|---|---|---|---|
| 0 | 0.04 | 0.80 (0.78 — 0.81) | 0.74 (0.72 — 0.75) |
| 1 | 0.04 | 0.80 (0.78 — 0.81) | 0.74 (0.72 — 0.76) |
| 2 | 0.04 | 0.80 (0.78 — 0.81) | 0.74 (0.72 — 0.76) |
| 3 | 0.04 | 0.81 (0.79 — 0.83) | 0.75 (0.73 — 0.77) |
| 4 | 0.04 | 0.81 (0.79 — 0.82) | 0.74 (0.72 — 0.75) |

**Myocardial infarction**

| $t_P$ | $I$ | Gradient Boosting AUROC | Reference AUROC |
|---|---|---|---|
| 0 | 0.03 | 0.71 (0.69 — 0.73) | 0.68 (0.65 — 0.70) |
| 1 | 0.03 | 0.73 (0.71 — 0.75) | 0.68 (0.66 — 0.71) |
| 2 | 0.03 | 0.74 (0.71 — 0.76) | 0.69 (0.67 — 0.71) |
| 3 | 0.03 | 0.72 (0.70 — 0.75) | 0.69 (0.67 — 0.72) |
| 4 | 0.03 | 0.73 (0.70 — 0.75) | 0.68 (0.66 — 0.71) |

**Stroke**

| $t_P$ | $I$ | Gradient Boosting AUROC | Reference AUROC |
|---|---|---|---|
| 0 | 0.03 | 0.72 (0.70 — 0.74) | 0.71 (0.69 — 0.73) |
| 1 | 0.03 | 0.75 (0.73 — 0.77) | 0.70 (0.68 — 0.72) |
| 2 | 0.03 | 0.75 (0.73 — 0.77) | 0.71 (0.69 — 0.73) |
| 3 | 0.04 | 0.75 (0.73 — 0.77) | 0.72 (0.70 — 0.74) |
| 4 | 0.04 | 0.75 (0.73 — 0.77) | 0.72 (0.70 — 0.74) |

**Cardiovascular disease**

| $t_P$ | $I$ | Gradient Boosting AUROC | Reference AUROC |
|---|---|---|---|
| 0 | 0.06 | 0.70 (0.69 — 0.72) | 0.64 (0.62 — 0.65) |
| 1 | 0.06 | 0.71 (0.70 — 0.73) | 0.64 (0.62 — 0.65) |
| 2 | 0.06 | 0.72 (0.71 — 0.74) | 0.65 (0.64 — 0.67) |
| 3 | 0.06 | 0.72 (0.70 — 0.73) | 0.65 (0.63 — 0.67) |
| 4 | 0.06 | 0.71 (0.69 — 0.73) | 0.64 (0.62 — 0.66) |

**Chronic kidney disease**

| $t_P$ | $I$ | Gradient Boosting AUROC | Reference AUROC |
|---|---|---|---|
| 0 | 0.03 | 0.77 (0.76 — 0.79) | 0.71 (0.69 — 0.73) |
| 1 | 0.03 | 0.77 (0.75 — 0.78) | 0.68 (0.66 — 0.70) |
| 2 | 0.03 | 0.77 (0.75 — 0.79) | 0.70 (0.68 — 0.72) |
| 3 | 0.04 | 0.79 (0.77 — 0.80) | 0.72 (0.70 — 0.74) |
| 4 | 0.04 | 0.79 (0.77 — 0.81) | 0.70 (0.68 — 0.72) |

**All-cause mortality**

| $t_P$ | $I$ | Gradient Boosting AUROC | Reference AUROC |
|---|---|---|---|
| 0 | 0.14 | 0.87 (0.86 — 0.87) | 0.80 (0.79 — 0.81) |
| 1 | 0.14 | 0.87 (0.87 — 0.88) | 0.81 (0.80 — 0.82) |
| 2 | 0.14 | 0.88 (0.87 — 0.88) | 0.81 (0.80 — 0.82) |
| 3 | 0.15 | 0.88 (0.87 — 0.88) | 0.82 (0.81 — 0.83) |
| 4 | 0.16 | 0.87 (0.87 — 0.88) | 0.80 (0.79 — 0.81) |

**Supplementary Table 5.** Prediction performance at one, two, three and four years after individual's first type 2 diabetes diagnosis. $t_P$, time of prediction counted in years since individual's first type 2 diabetes diagnosis; $I$, outcome incidence within study population; AUROC, area under receiver operating characteristic curve which confidence intervals were obtained by bootstrap sampling procedure.

| Logistic regression (LR) | |
|---|---|
| penalty | L2 |
| solver | lbfgs |
| C | 0.6, 0.7, 0.8 |
| max_iter | 300, 400, 500 |

| Random forest (RF) | |
|---|---|
| max_features | sqrt |
| criterion | gini |
| n_estimators | 1000, 1500 |
| max_depth | 12, 14, 16 |

| Gradient boosting (GB) | |
|---|---|
| n_estimators | 200 |
| booster | gbtree |
| eval_metric | auc |
| learning_rate | 0.025, 0.05, 0.1 |
| max_depth | 2, 3, 4, 5 |

**Supplementary Table 6.** Overview of model hyperparameters that were evaluated. For each model type, all combinations of listed hyperparameters were tested to identify those that led to the best average AUROC using 3-fold cross validation on the training set. Parameter names listed correspond to naming in software implementation (Python modules scikit-learn for LR and RF and xgboost for GB). Due to class imbalance (difference between number of cases and non-cases in each population) training error for class representing cases was scaled up by the inverse proportion between cases and non-cases.

**Heart failure ($I_{ts}$: 0.04, $I_{no-ts}$: 0.05)**

| | $AUROC_{ts}$ | $AUROC_{no-ts}$ |
|---|---|---|
| Reference, logistic regression | 0.74 (0.72 — 0.75) | 0.76 (0.74 — 0.77) |
| Logistic regression | 0.77 (0.76 — 0.79) | 0.78 (0.77 — 0.80) |
| Random forest | 0.77 (0.75 — 0.78) | 0.80 (0.79 — 0.81) |
| Gradient boosting | 0.80 (0.78 — 0.81) | 0.81 (0.80 — 0.83) |

**Myocardial infarction ($I_{ts}$: 0.02, $I_{no-ts}$: 0.03)**

| | $AUROC_{ts}$ | $AUROC_{no-ts}$ |
|---|---|---|
| Reference, logistic regression | 0.68 (0.65 — 0.70) | 0.69 (0.67 — 0.71) |
| Logistic regression | 0.70 (0.68 — 0.73) | 0.71 (0.69 — 0.73) |
| Random forest | 0.67 (0.64 — 0.69) | 0.71 (0.69 — 0.73) |
| Gradient boosting | 0.71 (0.69 — 0.73) | 0.72 (0.70 — 0.74) |

**Stroke ($I_{ts}$: 0.03, $I_{no-ts}$: 0.04)**

| | $AUROC_{ts}$ | $AUROC_{no-ts}$ |
|---|---|---|
| Reference, logistic regression | 0.71 (0.69 — 0.73) | 0.72 (0.70 — 0.73) |
| Logistic regression | 0.72 (0.70 — 0.74) | 0.72 (0.70 — 0.74) |
| Random forest | 0.69 (0.67 — 0.71) | 0.71 (0.69 — 0.73) |
| Gradient boosting | 0.72 (0.70 — 0.74) | 0.73 (0.72 — 0.75) |

**Cardiovascular disease ($I_{ts}$: 0.06, $I_{no-ts}$: 0.08)**

| | $AUROC_{ts}$ | $AUROC_{no-ts}$ |
|---|---|---|
| Reference, logistic regression | 0.64 (0.62 — 0.65) | 0.66 (0.64 — 0.67) |
| Logistic regression | 0.66 (0.65 — 0.68) | 0.68 (0.67 — 0.70) |
| Random forest | 0.67 (0.65 — 0.69) | 0.71 (0.69 — 0.72) |
| Gradient boosting | 0.70 (0.69 — 0.72) | 0.73 (0.71 — 0.74) |

**Chronic kidney disease ($I_{ts}$: 0.03, $I_{no-ts}$: 0.03)**

| | $AUROC_{ts}$ | $AUROC_{no-ts}$ |
|---|---|---|
| Reference, logistic regression | 0.71 (0.69 — 0.73) | 0.71 (0.69 — 0.73) |
| Logistic regression | 0.74 (0.72 — 0.76) | 0.75 (0.73 — 0.77) |
| Random forest | 0.74 (0.72 — 0.76) | 0.76 (0.74 — 0.78) |
| Gradient boosting | 0.77 (0.76 — 0.79) | 0.78 (0.76 — 0.80) |

**All-cause mortality ($I_{ts}$: 0.14, $I_{no-ts}$: 0.17)**

| | $AUROC_{ts}$ | $AUROC_{no-ts}$ |
|---|---|---|
| Reference, logistic regression | 0.80 (0.79 — 0.81) | 0.80 (0.79 — 0.81) |
| Logistic regression | 0.83 (0.82 — 0.83) | 0.83 (0.82 — 0.83) |
| Random forest | 0.85 (0.85 — 0.86) | 0.84 (0.84 — 0.85) |
| Gradient boosting | 0.87 (0.86 — 0.87) | 0.86 (0.86 — 0.87) |

**Supplementary Table 7.** Comparison of model performance between time-split and non-time-split models. AUROC measure for each prediction model types best parametrization (according to AUROC measure) for all outcomes compared between time-split ($AUROC_{ts}$; training, test and validation sets were split so that the model is trained on individuals diagnosed with type 2 diabetes historically earlier and evaluated on individuals diagnosed later) and non-time-split model ($AUROC_{no-ts}$; training, test and validation sets were split at random without accounting for date of prediction). $I_{ts}$ and $I_{no-ts}$, outcome incidence within time-split and non-time-split study populations, respectively. AUROC, area under receiver operating characteristic curve which confidence intervals were obtained through bootstrap sampling procedure.

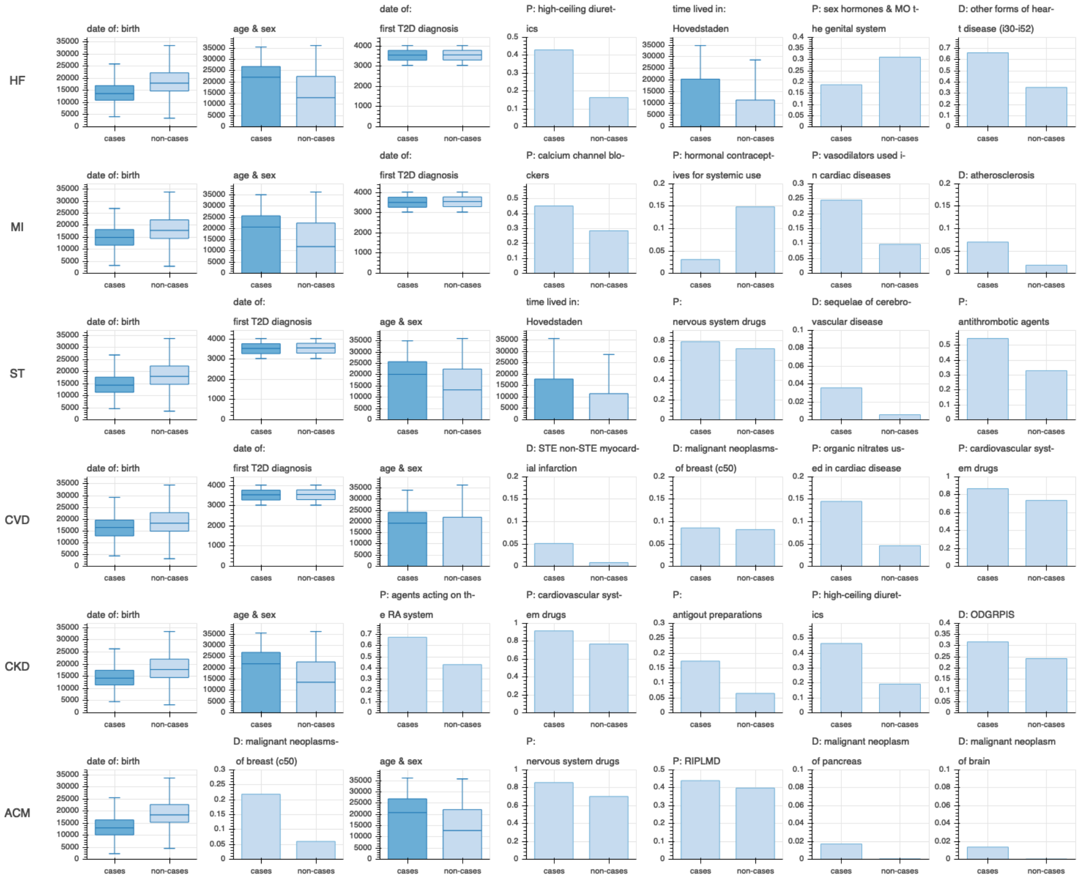


**Supplementary Figure 3.** Top seven most predictive gradient boosting features. For each comorbidity and ACM shown are the top seven features according to gradient boosting feature importance. Feature importance is an estimate of feature's relative contribution to outcome prediction. Box plots show a distribution of a given continuous feature (e.g. age, an interaction between age and sex) among cases and non-cases within validation set. Box plot whiskers represent lowest and highest observations still within 1.5 inter quantile range. To comply with Danish data protection rules, these values as well as values representing 25th, 50th and 75th percentiles were obtained by averaging five closest observations. Bar plots, describing count based features (e.g. count of a given drug prescription, count of a given diagnosis), show the proportion of validation set cases and non-cases with at least a single observation of that feature. HF, heart failure; MI, myocardial infarction; ST, stroke; CVD, cardiovascular disease; CKD, chronic kidney disease; ACM, all-cause mortality; D, diagnosis of; P, prescription of; MO, modulators of; STE, st elevation; RA, renin-angiotensin; ODGRPIS, other disorders of glucose regulation and pancreatic internal secretion; RIPLMD, hmg coa reductase inhibitors and plain lipid modifying drugs.

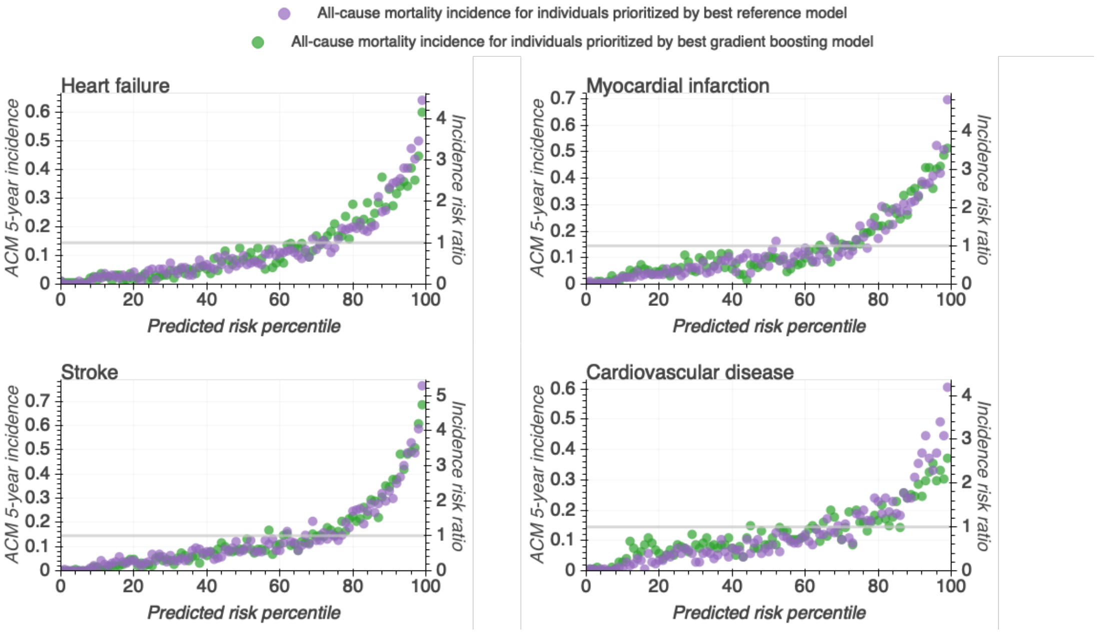


**Supplementary Figure 4.** All-cause mortality five-year incidence in population percentiles ranked by predicted type 2 diabetes comorbidity risk by the best gradient boosting (green) and best baseline (violet) models.
